# Ecological genomics of a novel host-parasitoid arms-race in nature

**DOI:** 10.64898/2026.06.30.735478

**Authors:** Leeban H. Yusuf, Jack G. Rayner, Renjie Zhang, Kalyani Z. Twyman, Michael Paulini, Xiao Zhang, Susan L. Balenger, Norman Lee, Robin M. Tinghitella, David A. Gray, Mark Blaxter, Nathan W. Bailey

**Affiliations:** University of St Andrews, Fife, Scotland; University of Maryland, Maryland, USA; Sanger Institute, Cambridgeshire, UK; Tianjin Normal University, Tianjin, China; University of Minnesota – Twin Cities, Twin Cities MN, USA; St. Olaf College, Northfield MN, USA; University of Denver, Denver CO, USA; California State University Northridge, Northridge CA, USA

## Abstract

Novel antagonistic interactions between species are expected to drive especially rapid coevolution. However, little is known about the genomic basis of such coevolution in nature because novel inter-specific interactions are rarely observed. Here, we study two species that recently came into first contact in Hawaii, the parasitoid fly *Ormia ochracea* and its cricket host *Teleogryllus oceanicus*. The fly locates crickets acoustically using their song, and parasitism usually results in host death. In response, protective male-silencing mutations have rapidly spread through cricket populations over the last ∼25 years, imposing novel selective pressure on flies. By integrating population genomic analyses of 358 re-sequenced flies with field surveys of selection imposed by host adaptations, we discover genomic signatures of recent selective sweeps driven by host adaptations, indicative of escalating arms-race dynamics. This evolutionary response is occurring despite severely depleted genetic variation after bottlenecks in Hawaiian fly populations. Comparative analyses suggest that the genomic substrate of modern-day, rapid counter-adaptation in *O. ochracea* has been under positive selection on intermediate and long-term timescales across parasitoid flies. Our findings thus support predictions of influential arms race coevolution models and illustrate the current and ancient genomic bases of counteradaptation in nature.

## Introduction

Human activity is driving novel antagonistic interactions between species, providing unprecedented opportunities to observe coevolution as it initiates (*1*, *2*). These interactions often lead to cycles of reciprocal adaptation that can shape species viability, diversity, and coexistence (*2–7*), yet the evolutionary mechanisms underlying reciprocal adaptation remain poorly understood. Classic theory predicts reciprocal adaptation should proceed via Red Queen dynamics, where frequency-dependent selection maintains balanced polymorphisms, or arms-race dynamics, where directional selection drives recurrent selective sweeps at coevolving loci (*8–10*). These two competing models generate distinct genomic signatures that allow researchers to identify the underlying dynamics of coevolution (*8*). However, empirical tests distinguishing these genomic dynamics remain rare and are confined largely to laboratory systems (*8*, *10*). Which, if either, of these models characterises novel coevolutionary dynamics in nature remains almost entirely unknown.

We studied this in *Ormia ochracea*, a crepuscular endoparasitoid fly found throughout the Americas and Caribbean that parasitises field crickets and katydids. The fly identifies hosts by localising host mating songs using a specialised auditory organ located in the fly’s prothorax (*11–13*). This hearing organ allows *O. ochracea* to pinpoint a sound source with incredible accuracy (<2° azimuth) (*14*). Upon host detection, gravid *O. ochracea* females deposit their larvae onto or around hosts, for whom infestation is usually fatal (*15–18*). In Hawaii, where *O. ochracea* has recently been introduced (first documented in 1989), it targets singing males of the field cricket *Teleogryllus oceanicus* as its preferred host (**Fig 1A & B**) (*19*). This interaction is unique since *O. ochracea* does not encounter *T. oceanicus* outside of Hawaii, and *T. oceanicus* is similarly not known to be parasitised by any acoustically orienting parasitoid throughout its native range in Australia (*20–22*). Under intense, recent parasitism by *O. ochracea* on male *T. oceanicus* in Hawaii, multiple, independent, adaptive male morphotypes have arisen in *T. oceanicus* populations and spread over the last 25 years, representing a textbook example of rapid adaptation due to novel species interactions (*23–27*). Each of these adaptive mutations disrupts sound-producing structures on cricket wings, protecting males from flies. Some of the mutations eliminate song completely, producing entirely silent male crickets (*20*, *28*); some provide protection by attenuating song (*29*, *30*); some generate novel signals that do not attract flies (*25*, *29*); others are attenuated in amplitude and bandwidth and avoid detection by flies yet remain detectable by, and attractive to, female crickets at close range (*25*, *31*). This suite of independent, adaptive mutations have recently spread to varying extents across a mosaic of Hawaiian cricket populations (**Fig 1D**) (*20*, *30*, *32–35*).

**Figure 1.**
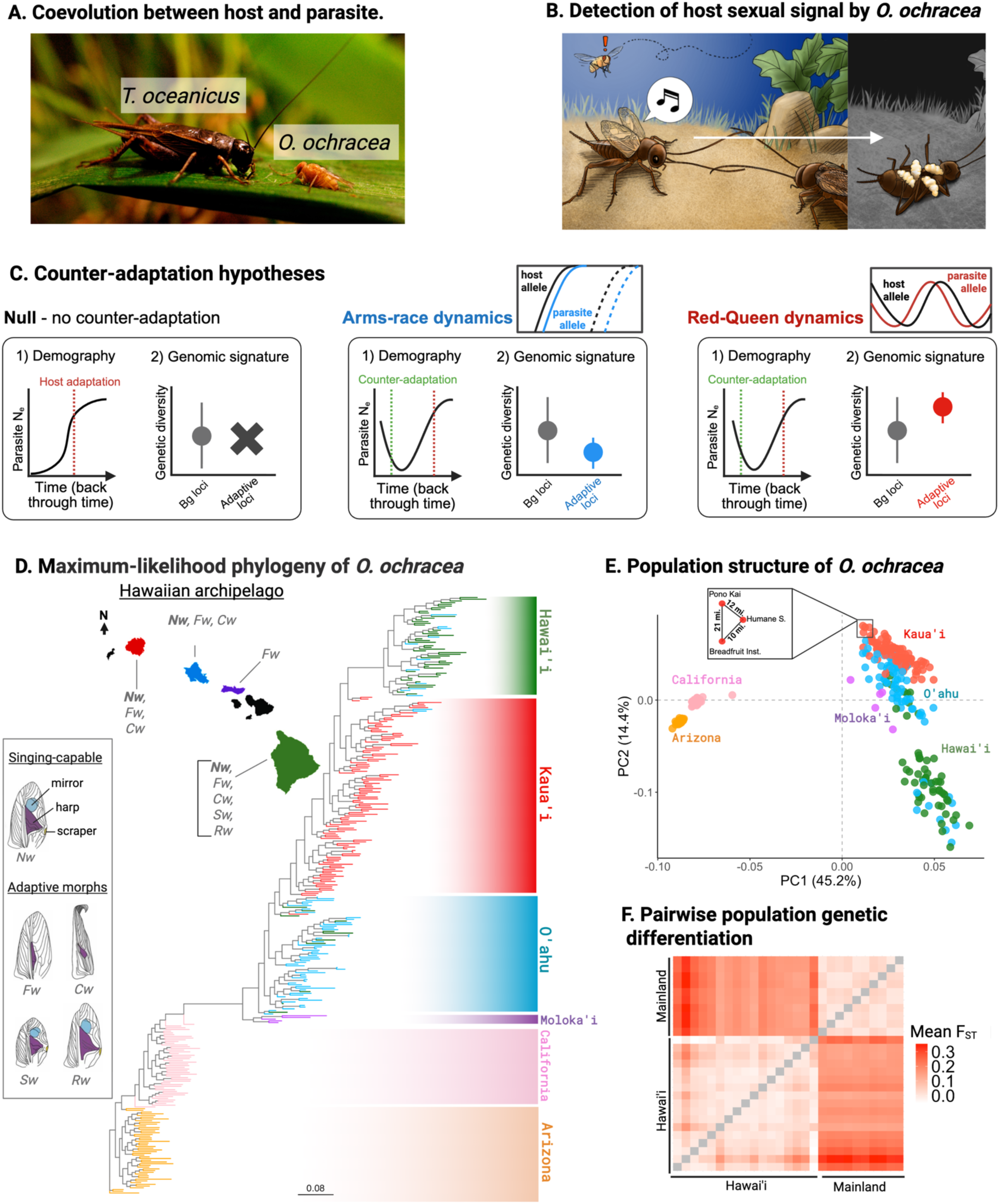
*Ormia ochracea and cricket coevolution*. **A.** *O. ochracea*, an eavesdropping parasitoid tachinid fly, parasitizes a novel, primary host (*T. oceanicus*) in Hawaii where it has been recently introduced. Figure credit and copyright: N. Lee. **B.** Upon detecting male advertisement song of *T. oceanicus*, *O. ochracea* deposits its planidia (live larvae) on or around singing males and burrows into them; larvae emerge 7-10 days later, killing the host. Figure credit: Anna Sieben, copyright: NW. Bailey. **C.** Antagonistic co-evolutionary models (arms-race and Red-Queen dynamics) and hypotheses of fly counter-adaptation. Assuming no counter-adaptation, we expect significant *O. ochracea* population declines and no genomic regions responding to selection due to recent host adaptation. If counter-adaptation is occurring under one or both models, we expect a recent *O. ochracea* population size recovery. At loci underpinning counter-adaptation, arms-race predicts a selective sweep characterised by local reduction in genetic diversity, while Red-Queen predicts a local increase in genetic diversity relative to background (’Bg’) loci. **D.** In Hawaii, adaptive host morphs (’flatwing’ (*Fw*); curly-wing (*Cw*), small-wing (*Sw*), rattling-wing (*Rw*)) have evolved attenuated sound-producing structures (mirror, harp and scraper) on their wings in response to selection imposed by *O. ochracea* (Figure adapted from (*19*)). Map shows the distribution of adaptive morphs across the Hawaiian archipelago. Little is known about the evolutionary dynamics of *O. ochracea*. Maximum-likelihood tree of 285 whole-genome resequenced *O. ochracea* collected across four islands in 2022. **E.** PCA of 347 whole-genome samples (285 collected in 2022; an additional 62 in 2024), coloured by sampling location. Box highlights four Kaua’i samples from distinct sampling locations, illustrating little to no local population structure within islands. **F.** Mean pairwise population differentiation (F_ST_) between Hawaiian and US mainland populations (Californian and Arizonan populations).

Fly larvae must infest hosts to complete development. Although other suitable host species of crickets occasionally appear in Hawaii, there is no evidence that other host species sustain Hawaiian *O. ochracea* populations (*16*, *36*). Therefore, the spread of adaptive *T. oceanicus* morphotypes, whose songs have shifted in amplitude or frequency relative to those *O. ochracea* has evolved to localise, should impose strong natural selection on Hawaiian *O. ochracea*. Field surveys across its American and Hawaiian range show that *O. ochracea* is locally adapted to the most common regional host, and electrophysiological recordings reveal that Hawaiian flies exhibit different neural responses to the sounds made by incompletely-silenced *T. oceanicus* morphs compared with ancestral populations (*21*, *37*). This system offers a rare window into the potential initiation of reciprocal adaptation, capturing coevolution as it begins to unfold during a novel interaction.

We investigated the genomic basis and evolutionary mechanism underlying counter-adaptation in *O. ochracea* to novel host adaptations in Hawaii (**Fig 1C**). We collected and sequenced (30xcoverage) 249 wild-caught *O. ochracea* individuals across 27 populations spanning four islands of the Hawaiian archipelago. We performed field surveys in 12 focal *T. oceanicus* populations to quantify host morphotypes and conducted gene expression experiments in gravid female flies to identify genes involved in host detection. To contrast any genomic signatures of recent coevolution with those arising over longer historical timeframes, we also sequenced 99 *O. ochracea* individuals from six populations in California and Arizona, where this fly parasitizes at least six different *Gryllus* species, and 10 individuals from the closely related outgroup *O. depleta*, introduced from South America to Florida in the 1980s-90s for biological pest control (*38*). Across mainland populations of *O. ochracea*, their cricket hosts show no evidence of adaptation due to parasitism via attenuation of male calling song. Using this population genomic dataset, we inferred when *O. ochracea* arrived in Hawaii, characterized population structure, and tested for genomic signatures of selection. Our results reveal that multiple selective sweeps underlie rapid counter-adaptation in *O. ochracea*, providing genomic evidence for arms-race dynamics in natural coevolving populations. Finally, comparative genomic analysis of the parasitoid fly family *Tachinidae* (*16*) shows that genes underlying recent coevolution have also experienced a much longer evolutionary history of positive selection, implicating the recruitment of ancestrally-selected genes in rapid evolutionary responses to novel host-parasite encounters.

## Results

### A novel host-parasite interaction: *Ormia ochracea* recently arrived in Hawaii

We examined evolutionary relationships among *O. ochracea* populations across Hawaii and outgroup populations using principal component analysis (PCA) and maximum-likelihood (ML) phylogenetic trees. Hawaiian populations formed a single clade and generally clustered by island (**Fig 1D** and **E**). In both PCA and ML trees, individuals showed little grouping by sampling location, suggesting frequent migration within islands (**Fig 1E**). Some individuals from O‘ahu and Hawai‘i (Big Island) clustered together, indicating bidirectional inter-island gene flow, a pattern also seen in *T. oceanicus* (*32*, *34*). Arizonan and Californian *O. ochracea* populations also showed little genetic differentiation between sampling localities or individuals collected using different host song playbacks (**Fig 1F** and **Supplementary Fig. 1**). Overall, genetic differentiation was slightly higher among Hawaiian populations (mean F_ST_ = 0.024) than among North American ones (mean F_ST_ = 0.018), with moderate differentiation between Hawaiian and North American populations (mean F_ST_ = 0.188) (**Fig 1F**). Within islands, differentiation was low on Kaua‘i and Hawai‘i (mean F_ST_ ≤ 0.001) but higher on O‘ahu (mean F_ST_ = 0.024), suggesting older or more structured populations there. Collectively, these results indicate metapopulation dynamics in *O. ochracea*, with weak but detectable structuring between Hawaiian islands and ongoing inter-island gene flow.

Using demographic modelling of *O. ochracea* from Hawaii (n=186) and North American (n=99) samples collected in the same year, we show that the fly is likely to have arrived in Hawaii ∼50 years ago (600 generations, generation time of 1-2 months (*39*)), and that since then there has been bidirectional gene flow between North American and Hawaiian populations (**Fig 2A**). Historical effective population size modelling using a Bayesian implementation of PSMC demonstrates a population crash in Hawaiian flies between 100 – 1,000 generations ago (**Fig 2B**), around the inferred split time of Hawaiian populations from North American populations, and suggests that *O. ochracea* arrived shortly before their first identification in Hawaii in 1989 (*40*). These results place a contemporary, recent upper limit on the initiation of this coevolutionary interaction, meaning any genomic and phenotypic signatures of adaptation must have emerged within decades of first contact between the parasitoid and its host.

**Figure 2.**
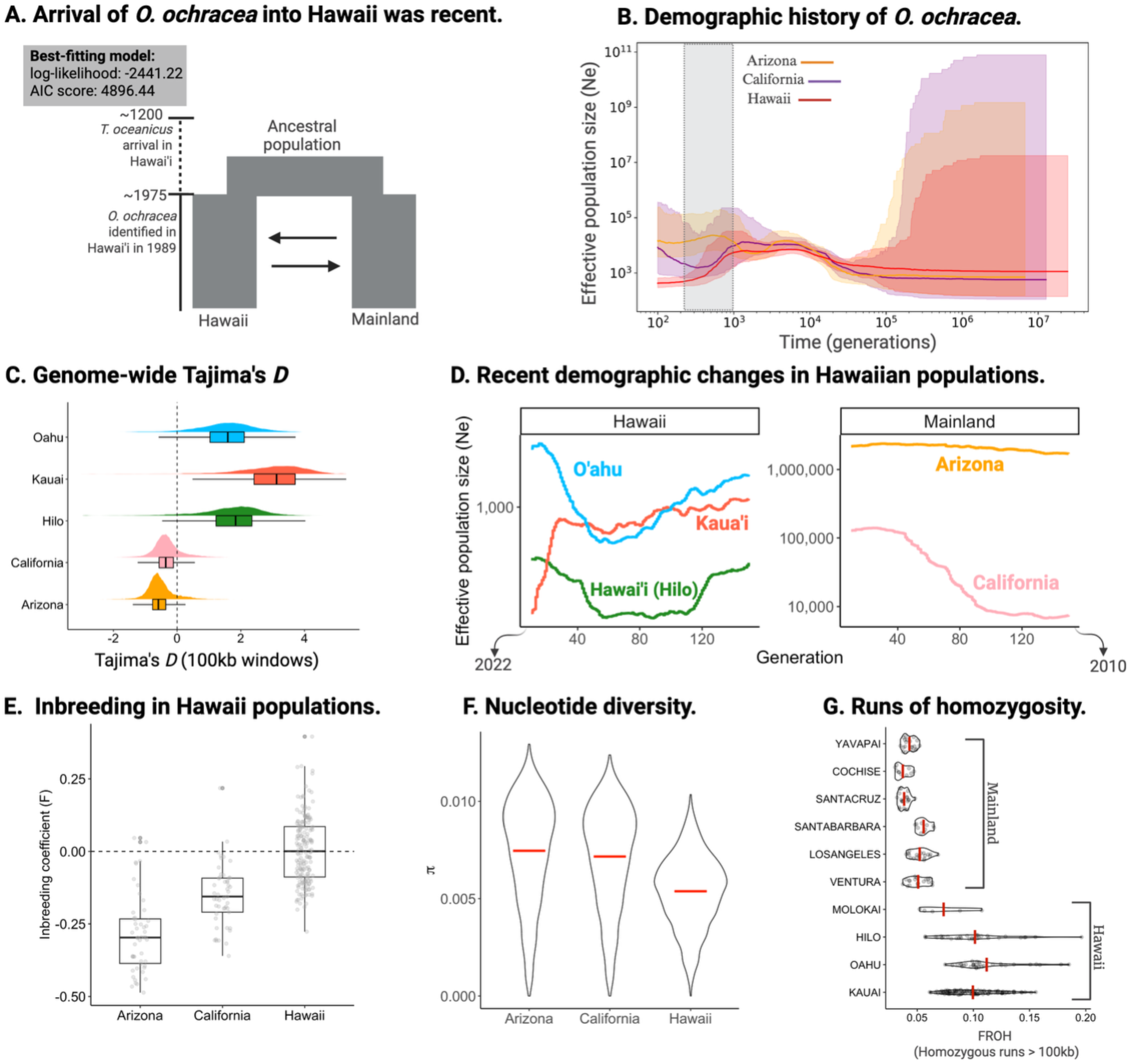
**Evolutionary history of Hawaiian *Ormia ochracea. A.*** Best-fitting demographic history of the split between Hawaiian and US mainland (Californian and Arizonan) populations of *O. ochracea.* Based on previous analyses (Zhang et al., 2020), *T. oceanicus* arrived with Polynesian settlers ∼800 years ago. According to the best-fitting demographic history (using *moments* in *GADMA2*) (*91*, *92*), *O. ochracea* arrived ∼50 years ago, closely matching initial observations of *O. ochracea* in Hawaii in 1989. ***B.*** Effective population size through time (in generations) based on coalescent analysis using whole-genome samples of *O. ochracea* from Hawaii, California and Arizona. Grey rectangle roughly illustrates divergence between Hawaiian and US mainland samples inferred by *moments*, coinciding with a sustained drop in effective population size in Hawaiian samples. ***C.*** Tajima’s *D* calculated in 100kb windows for whole-genome samples of *O. ochracea* from Hawaiian Islands and the mainland. Dashed line indicates neutral expectation (similar observed and expected genetic variation)*. **D.*** Linkage-based analysis of recent (∼10 years) effective population size changes using island-specific whole-genome samples of *O. ochracea. **E.*** Inbreeding coefficient in Hawaiian, Californian and Arizonan samples *of O. ochracea.* Dashed line indicates observed heterozygosity matches expected heterozygosity. Values below the dashed line indicate excess heterozygosity and above indicates reduced heterozygosity*. **F***. Nucleotide diversity in Hawaiian, Californian and Arizonan samples of *O. ochracea. **G.*** Fraction of runs of homozygosity (considering only homozygous runs more than 100 kb) for Hawaiian and US mainland samples. Larger FROH indicate a larger proportion of the genome is found in a run of homozygosity.

Estimates of Tajima’s *D* provide further evidence for a recent introduction in Hawaiian populations of *O. ochracea* (**Fig 2C**). Californian and Arizonan populations show levels of genome-wide polymorphism consistent with a neutral equilibrium, whereas across Hawaiian populations we observed extreme genome-wide positive Tajima’s *D,* consistent with a very recent population contraction, and little support for population recovery (*41*, *42*) (**Fig 2C**). To corroborate population dynamics inferred from genome-wide Tajima’s *D* estimates, we estimated recent demographic changes in populations of *O. ochracea* across Hawaii using genome-wide patterns of linkage disequilibrium. Island-level estimates of N_e_ through time over the last 150 generations from our focal sampling point in 2022 show *O. ochracea* populations have extremely low levels of N_e_, with estimates across islands ranging from ∼500 - 1400. We observe island-specific fluctuations in N_e_, with recent crashes observed across all three islands, and at the same time periods (100 - 40 generations ago). Parallel crashes in recent N_e_ have been observed in *T. oceanicus* (*34*), suggesting recent population fluctuations in *O. ochracea* may be mediated by similar population dynamics in *T. oceanicus*. *O. ochracea* populations across Hawaii appear to have arrived in Hawaii experiencing a strong founder effect with little evidence of population recovery, potentially due to host adaptations evolving as early as ∼2003 (*20*, *23*). As a result of these demographic circumstances, inbreeding is higher in Hawaii compared to North American populations (**Fig 2E**). Additionally, genetic diversity is lower in Hawaiian populations compared with North American populations (**Fig 2F**), and the fraction of the genome in runs of homozygosity (FROH) is higher across samples collected in Hawaii (**Fig 2G**).

### Selective sweeps in Hawaiian *O. ochracea* are associated with host-seeking behaviour

Whether arms-race or Red Queen dynamics explain rapid coevolution between novel interactants is an open question. Arms-race models predict repeated fixation of beneficial mutations via selective sweeps, whereas Red Queen dynamics predict the maintenance of genetic variation at coevolving loci by balancing selection. Adaptive signal-attenuating mutations in cricket hosts have rapidly risen to high frequency across Hawaii, and in some cases have fixed (*32*, *43*). We therefore hypothesised that counter-adaptation in Hawaiian *O. ochracea* populations might therefore be characterised by similar dynamics. However, population crashes and bottlenecks can also have pronounced effects on genome-wide variation, which requires us to disentangle demography from selection in signatures of genomic variation.

First, we computed three statistics to identify potential selective sweeps in Hawaiian populations of *O. ochracea*: a (1) likelihood-based approach (SweeD) to detect selective sweep signatures using phased whole-genomes (*44*); (2) the normalized population branch statistic (*PBSn1*) which detects local genetic differentiation in a focal population (Hawaii)(*45*); and (3) the cross-population haplotype statistic (XP-EHH) (*46*), which detects recent positive selection by comparing haplotype homozygosity between Hawaiian and mainland populations.

We identified 67 genomic outlier windows on the second, third and fourth autosomes, which were consistently represented in the top 5% of the distribution across all three statistics (**Fig 3A and B**). These outlier windows cluster in three narrow genomic regions, one on each chromosome, spanning from 1.5 Mb – 7.8 Mb and consisting of 14 – 26 protein-coding genes. Alongside sweep scans, McDonald-Kreitman tests reveal stronger patterns of adaptive evolution genome-wide and within these sweeps in Hawaiian populations of *O. ochracea* relative to the US Mainland (**Supplementary text S1; Supplementary Fig. 3**). Protein-coding genes within these 67 outlier windows were enriched for biological and molecular ontology categories involved in ionotropic glutamate receptor signalling pathway (*P*_adj_ = 0.008) and glutamate-gated calcium ion channel activity (*P*_adj_ = 0.001). Genes involved in glutamate signalling were found in each candidate sweep region and include: *GRIK1* (Glutamate receptor, ionotropic, kainate 1), an ionotropic receptor involved in synaptic function and neurotransmission (*47*); *APP* (Amyloid beta precursor protein), a transmembrane protein involved in synapse formation and plasticity (*48*, *49*); and *GRIN2B* (Glutamate receptor subunit epsilon-2), which encodes a subunit of a glutamate receptor and is involved in excitatory neurotransmission, synaptic plasticity, memory and learning (*50*) (**Fig 3D**).

**Figure 3:**
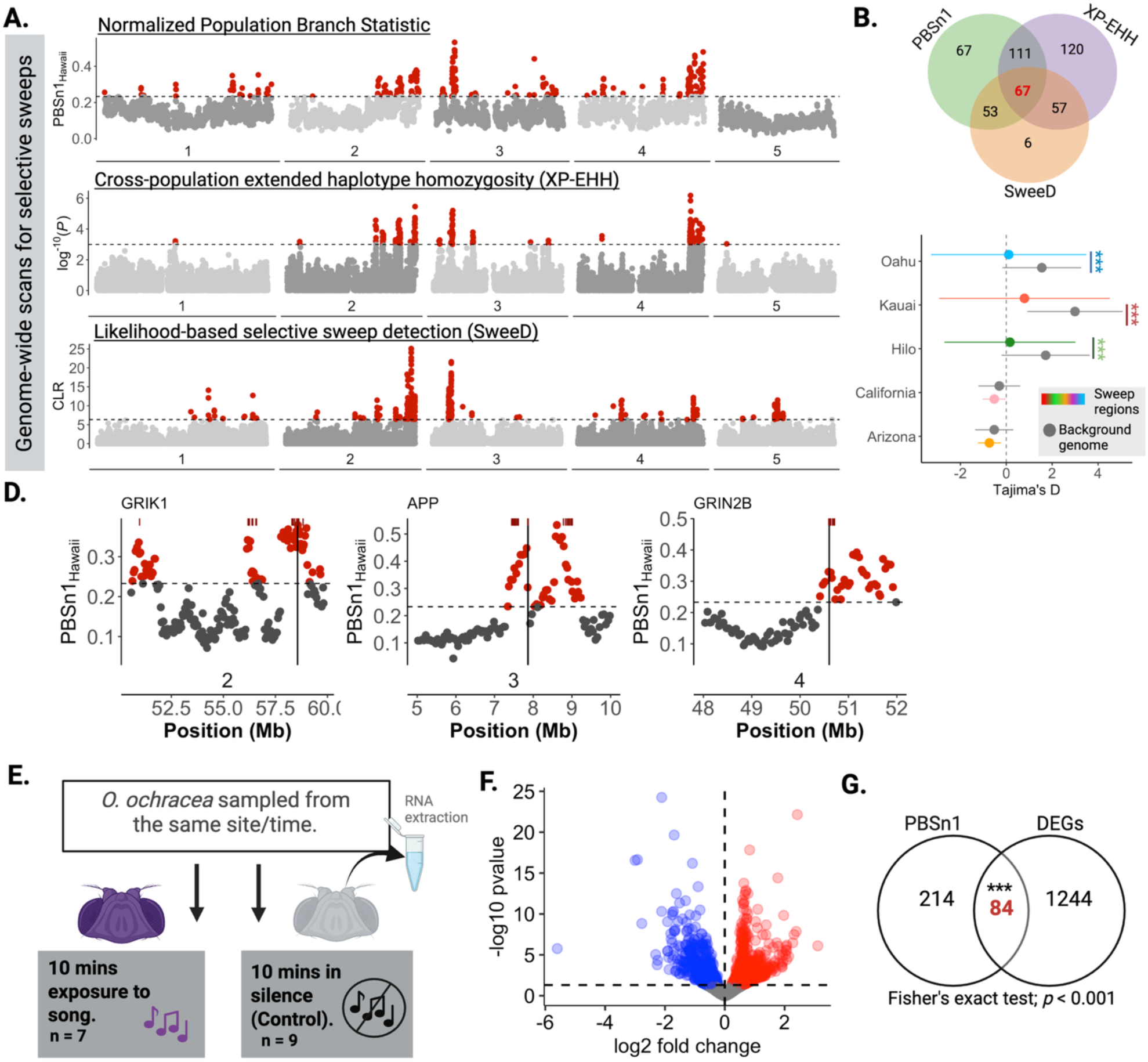
Evidence of adaptation by directional and potentially balancing selection on *O. ochracea.* **A.** Genome-wide selection scans across the five *O. ochracea* chromosomes for selective sweeps including PBSn1_Hawaii_ (population branch statistic comparing the differentiation in the focal population (Hawaii) relative to differentiation between Californian and Arizonan populations, rescaled by the total length of the three-population tree), XP-EHH (FDR adjusted p-value) and CLR likelihood-test implemented in SweeD. The dashed line indicates the 95% quantile and red dots are outlier windows. **B.** Overlap between outlier regions across all three selective sweep scans. **C.** Tajima’s *D* for candidate selection sweep regions (n=67) compared to the background genome across Hawaiian islands and in North American populations. Point represents mean Tajima’s *D* per island/state, and bars indicate ±1 SD. Stars indicate level of significance: 0.0001 = ***, 0.001 = **, 0.01 = *, 0.05 =., >0.05 = ns in Mann-Whitney U test. **D.** Normalised Population Branch Statistic (PBSn1_Hawaii_) at three candidate loci under directional selection in Hawaii (*GRIK1, APP* and *GRIN2B*). Dashed line indicates the 95% quantile and red dots indicate outlier windows. Positions of protein-coding genes shown in maroon rectangles above plot and solid vertical line indicates the positions of example candidate genes. **E.** Field-based RNA-seq to detect genes involved in host song detection and perception in *O. ochracea*. **F.** Volcano plot; dots above the horizontal dashed line indicate genes with significant differential expression (*p*<0.05 after BH correction). Genes upregulated in *O. ochracea* female heads in response to *T. oceanicus* song are shown in red; downregulated genes are shown in blue. **G.** Venn diagram showing overlap between DE genes and genomic outlier windows identified in PBSn1 scan for selective sweeps in Hawaii. Stars indicate level of significance: 0.0001 = ***, 0.001 = **, 0.01 = *, 0.05 =., >0.05 = ns in Fisher’s Exact test.

To rule out potential confounding effects of demographic history on our selective sweep inference, we used SLiM (v. 4.3) (*51*) to simulate a severe and recent population bottleneck in Hawaiian *O. ochracea*, and re-computed likelihood-based selective sweep summary statistic (SweeD) using simulated genetic variation. Using the highest composite likelihood ratio (CLR = 6.34) identified in the simulation as a cut-off in our observed data, we still recover the three major outlier regions, indicating that demography alone is unlikely to putative selective sweeps (**Supplementary Fig. 2**). Additionally, we assessed whether selective sweeps are occurring across islands rather than reflecting a strong signal in a single island. We compared Tajima’s *D* in sweep regions to the background genome across islands. Tajima’s *D* was consistently lower in sweep regions than in the background genome across all three islands (*Mann-Whitney U* test, all tests *p* < 0.001): Kaua’i (sweep: −0.79, background: 2.99), O’ahu (sweep: 0.10, background: 1.55), and Hawai’i (sweep: 0.16, background: 1.72) (**Fig. 3C**). Together, these results imply that strong directional selection is acting in Hawaiian populations of *O. ochracea*.

Given that signals of rapid directional selection are acting on genes that may influence behaviour, and that attenuation of host mating signals is an important selective pressure on *O. ochracea* populations in Hawaii, we asked whether genes under selection in Hawaiian populations are involved in host-seeking and detection (**Fig 3E**). We identified differentially expressed genes (DEGs) in the heads of gravid *O. ochracea* females exposed to *T. oceanicus* song compared to those not exposed. We found 1,328 genes (out of 11,603 tested genes) which showed (P_adj_ < 0.05) differential gene expression in gravid female brains during exposure to *T. oceanicus* song (**Fig 3F**). Of these 1,328 DEGs, 638 were downregulated and 690 were upregulated in response to *T. oceanicus* song. These genes appear to be involved in cytoplasmic processes (*P*_adj_ < 0.001) and small molecule metabolism (*P*_adj_ < 0.001). We asked whether genes within sweeps are overrepresented in DEGs, consistent with a link between directional selection and host-seeking behaviour. We found a significant overlap between sweep outlier regions identified using the *PBSn1* statistic (n= 298; *PBSn1* > 0.232) and differentially expressed genes (n=1,328) (**Fig 3G**) (Fisher’s exact test; *p* < 0.001). We found similar significant overlap between DEGs and sweep outlier regions identified by SweeD (Fisher’s exact test; *p* < 0.001) and XP-EHH (Fisher’s exact test; *p* < 0.001). Only 8 genes were both significant across all sweep tests and differentially expressed in gravid female heads in response to *T. oceanicus* song. Among these 8 overlapping genes, located across the second, third, and fourth autosomes, three are of particular interest: *GRIK1* (described above), *MAP1B/futsch*, a microtubule protein involved in axonal growth and synaptogenesis (*52*), and *TSC22D/bunched*, a transcription factor linked to mushroom body development in *Drosophila* (*53*). These results imply that directional selection in Hawaiian *O. ochracea* is associated with a functionally diverse subset of genes that are down-regulated in heads of Hawaiian *O. ochracea* during exposure to host sexual signals.

### Genomic evidence of recent counter-adaptation in *O. ochracea*

Our last field sampling effort (June-Oct 2022) demonstrates the considerable variation in proportions of normally-singing versus adaptive cricket morphs across sites and islands in Hawaii, with some populations consisting of mostly normally-singing males (e.g. Kauai-PK: 73%) and others containing none (Kauai-AS: 0%) (**Fig 4A** and **Supplementary Table 1**). The proportion of adaptive morphs observed across populations at our final sampling point is similar within years (2022 January and June), but we observe a gradual decrease in singing-capable males across years (**Fig 4A**), presumably due to fly parasitism and the emergence of multiple adaptive host morphs. We used this field survey data to test whether *O. ochracea* populations in Hawaii are responding to *T. oceanicus* adaptive morphs. We analysed the association between genetic variation across 12 *O. ochracea* populations (n=151 individuals) with the proportion of singing-capable *T. oceanicus* males surveyed from those same populations in a latent factor mixed model (*LFMM*).

**Figure 4.**
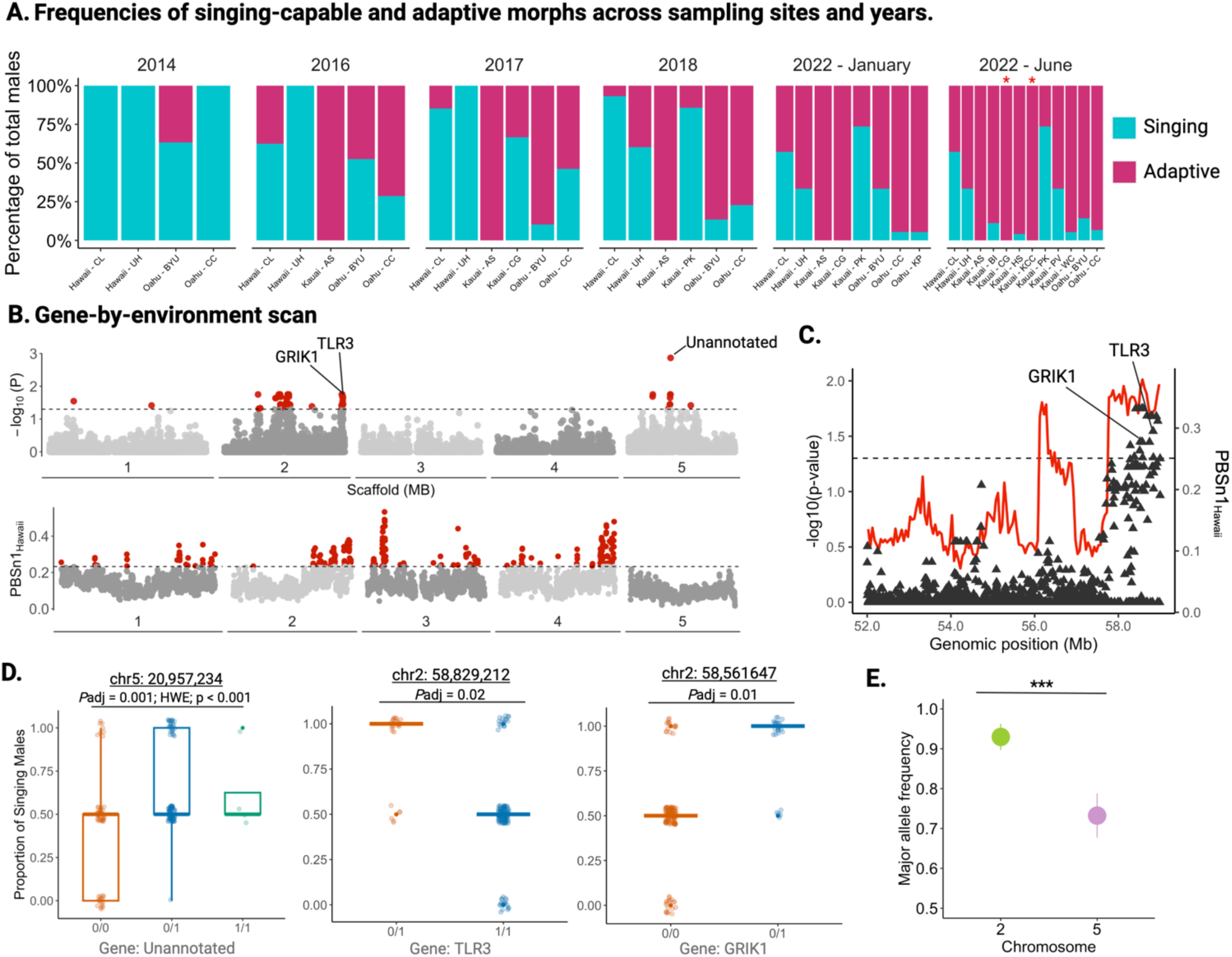
Associations with the host adaptation environment reveal genomic signatures of fly counteradaptation. **A.** Frequencies of singing-capable males and adaptive male morphs of *T. oceanicus* across sites in Hawaii. Red asterisks denote populations where singing-capable males were heard during sampling but not collected. **B.** Gene-by-environment scan across the five chromosomes of *T. oceanicus* using the proportion of singing-capable males identified in 12 Hawaiian focal populations as a predictor of genetic variation. We identified two main regions (chromosome 2 and 5) with significant associations to frequencies of singing-capable morphs across populations in Hawaii. Dashed line and red points above it (BH-adjusted *p* < 0.05) indicate significance level and significantly-associated SNPs, respectively. Below the first genome-wide Manhattan plot, is a selective sweep scan (PBSn1) for comparison. **C.** Region on chromosome 2 with strong support for rapid directional selection and counter-adaptation. Black triangular points indicate SNPs in gene-by-environment (GEA) scan and dashed line (BH-adjusted *p* < 0.05) indicates significance and corresponds to the left axis, and red line indicates PBSn1 values and correspond to right axis. **D.** Boxplots show genotypes for examples of associated SNPs within or close to (<1000 bp) candidate genes in GEA analyses. The center line in the boxplots represent the median proportion of singing-capable males per genotype and whiskers are 1.5 x interquartile range, with individuals represented as points. **E.** Mean major allele frequency (± 1 SD) for SNPs identified in GEA analyses on chromosomes 2 and 5. Stars indicate level of significance: 0.0001 = ***, 0.001 = **, 0.01 = *, 0.05 =., >0.05 = ns in Mann-Whitney U test.

This analysis yielded 67 variants, concentrated in two separate genomic regions on the second and fifth autosomes, that predict the proportion of singing-capable males (**Fig 4B** and **Supplementary Table 2**) (*P*_adj_ < 0.05). The most highly associated SNP (chr5: 20,957,234) showed significant departure from Hardy-Weinberg equilibrium (HWE) because of heterozygote excess (χ^2^ = 19.52, *p* < 0.001) (**Fig 4B** and **4D**). In fact, of the 11 significantly associated variants found on the fifth autosome, nine show significant departures from HWE due to heterozygote excess and span almost a 1 Mb genomic region **(Supplementary Table 2** and **3)**. For significantly-associated variants on the fifth autosome, heterozygous individuals often persist better in populations consisting of high/intermediate (e.g. chr5: 20,957,234; n=26) and low (n=63) proportions of singing-capable *T. oceanicus*, whereas individuals homozygous for the reference allele are less commonly found in high/intermediate song environments (n=8), but are increasingly able to persist in low song (n=32) and ‘silent’ host populations (n=17) (**Fig 4D**).

In contrast, only two SNPs outside of the fifth autosome show significant departures from HWE, with most other significant SNPs characterised by a deficit of heterozygotes **(Supplementary Table 3)**. Variants significantly associated with the proportion of singing-capable males on the second autosome, for example, show allele frequencies that almost reach fixation at the major allele (**Fig 4E**). Our results suggest genomic counter-adaptation may be occurring by two contrasting genomic mechanisms owing to two different selective modes: pervasive selective sweeps driven by directional selection, and rarer heterozygote advantage driven by recent balancing selection. Outside of Hawaii, we found little convincing evidence for balanced polymorphisms in demographically stable Arizonan and Californian populations of *O. ochracea*, where genomic tests of long-term balancing selection are more viable (**Supplementary text S2; Supplementary Fig. 4**).

Of the 67 significant variants, 22 are found within 20 protein-coding genes (10 with annotations), 3 are within two long non-coding RNAs, and the 42 remaining variants are intergenic. Three genes (*GRIK1*, *TLR3*/*Toll-7* and an uncharacterised protein) contain variants associated with the proportion of singing-capable males across Hawaii and are outliers in all three selective sweep statistics (**Fig 4B**). *TLR3/Toll-7* and *GRIK1* are both found near the end of the second autosome and encode a toll-like receptor involved in antiviral defence (TLR3/Toll-7), as well as neutrophin signalling and odorant receptor function (*54–56*), and a glutamate receptor (*GRIK1*), which we described above (**Fig 4C**). Of particular interest is *GRIK1*, which is under strong directional selection, is significantly associated with the proportion of singing-capable males across Hawaii, and is differentially expressed in female *O. ochracea* heads when exposed to *T. oceanicus* song. Inference of the strength of selection on both *GRIK1* and *TLR3* using approximate full-likelihood estimates of selection coefficients (*CLUES2*) suggests that directional selection (*s* = 0.028 – 0.038) at these loci is incredibly strong and intensified recently (**Supplementary text S3; Supplementary Fig. 5; Supplementary Table 4**). Several intergenic variants found to be significantly associated with the proportion of singing-capable males across Hawaii flank these genes, particularly *TLR3/Toll-7*, implying that these associated variants may impose functional changes in both protein structure and gene expression linked to counter-adaptation in *O. ochracea* (**Supplementary Table 2**).

### Fly counteradaptation genes are associated with Tachinid diversification

Adaptation to detect and specialize on novel hosts has been documented across *O. ochracea’s* range (*21*, *22*), and following its introduction to Hawaii (*37*). This local specialisation may have facilitated the spread of *O. ochracea*, but also divergence of spatially isolated *O. ochracea* populations, or species within the genus. We investigated the evolutionary lability of host choice, and its effect on *O. ochracea* divergence, by sequencing 10 individuals from a closely related species, *O. depleta*, which is found in South America but was introduced to North America in the 1990s to control mole cricket populations (**Fig. 5A**) (*38*). Beyond divergence in host preference, little else appears to differentiate North American populations of *O. ochracea* from *O. depleta*.

**Figure 5:**
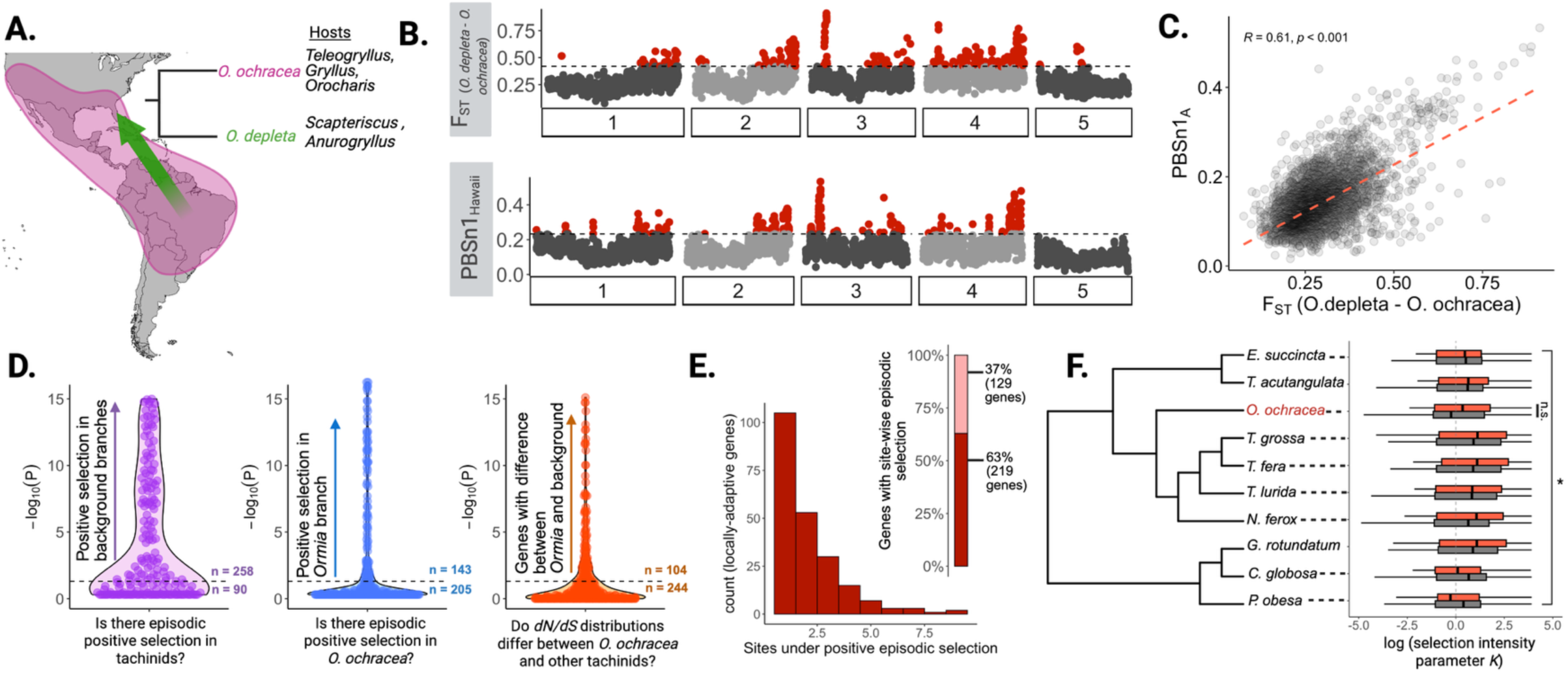
Historic episodic positive selection is a common feature of genes involved in recent fly adaptation to host defenses. **A.** Model of *O. ochracea* and *O. depleta* evolutionary history. Pink highlighting shows approximate distribution of *O. ochracea* throughout the Americas. *Ormia depleta* was introduced into Florida from Brazil to control mole cricket (*Scapteriscus* spp.) populations. **B** and **C.** Genetic differentiation (F_ST_) between *O. ochracea* and *O. depleta* is strongly associated with local genomic differentiation in Hawaiian *O. ochracea* (PBSn1). **D.** BUSTED-PH; gene-wide tests for episodic positive selection. Dashed lines indicate significance threshold (*p* < 0.05) and values above and below each violin plot indicate the number of locally adaptive genes (in sweep regions found in Hawaiian *O. ochracea*) that show evidence for episodic positive selection in each test and those that do not, respectively. Background branches refer to all branches, which include nine other tachinid species, in gene trees that are not *O. ochracea.* Most locally adaptive genes show evidence of positive selection in other tachinids. **E.** Histogram showing the number of sites (per gene) with evidence of episodic positive selection. Bar plot highlights the number of genes with at least one site under positive selection (dark red) across all ten tachinids tested including *O. ochracea*, and genes showing no evidence of site-wise positive selection (light red). **F.** Species tree of tachinids used in this study. RELAX tests show that genes involved in fly adaptations to host cricket defenses (red) show, on average, evidence of intensified selection (log (*K* > 1)) rather than relaxed selection (log (*K* < 1), compared to all other genes in the genome. The center line in the boxplots represent the median log (*K*) and whiskers are 1.5 x interquartile range for log(*K*), in genes under local selection in Hawaiian *O. ochracea* populations (red) and background genes (grey).

If arms-race dynamics drove divergence in host detection and specialisation between *O*. *ochracea* and *depleta*, we would expect genomic regions underlying counteradaptation to be associated with regions that differentiate these two species (*8*). In contrast, Red-Queen dynamics are characterised by balancing selection, which is expected to maintain local polymorphism and minimise genomic divergence at loci involved in coevolution (*57*). We detected genome-wide differentiation (F_ST_ computed in 50 Kb windows) between *O. ochracea* on the mainland and *O. depleta* from Florida, and found that regions of extreme F_ST_ strongly overlap with candidate selective sweeps specific to Hawaiian *O. ochracea* populations (**Fig. 5B**) (Fisher’s exact test; p < 0.001). Additionally, the degree of genetic differentiation specific to Hawaii (measured using *PBSn1*) was strongly correlated with genetic differentiation (F_ST_) between North American *O. ochracea* and *O. depleta* across the entire genome (Pearson’s correlation; 5,786 50kb windows, *r* = 0.61, *p* < 0.001) (**Fig. 5C**). Thus, the genomic basis of counteradaptation within *O. ochracea* involves the same genomic regions implicated in interspecific divergence with its sister species.

It is unknown whether genes coevolving during novel host-parasite interactions represent new targets of selection, or whether they are ‘usual suspects’ that recurrently experienced selection through deeper evolutionary time. We tested this using comparative genomic analyses of nine other tachinid parasitoids with high-quality genomes (**Supplementary text S4**; **Supplementary Table 5** and **Figs. 7 – 10).** Using a suite of phylogenetic tests, we first demonstrate with gene-wide analyses of episodic directional selection that the majority of orthologues within recent Hawaii-specific selective sweeps (n = 348) also experienced positive selection during the diversification of tachinids (*BUSTED-PH*). Of the 348 genes tested, 258 (likelihood ratio test; *p* < 0.05) showed evidence of positive episodic selection in tachinids on background branches (not including *O. ochracea*), with fewer (n=143) showing evidence of episodic positive selection in just *O. ochracea* (**Fig. 5D**). Site-wise tests (*FUBAR*) of positive selection indicate that 219 of these genes harbour at least one codon experiencing positive selection across parasitoid flies (posterior probability > 0.9) (**Fig. 5E**). Just under a third of all tested genes (n=104) differed in *dN/dS* between background branches and *O. ochracea,* implying selective regimes in this subset of genes recently shifted. Evidence from a complementary analysis testing for a change in selective regime (*RELAX*) show that this is unlikely to be due to relaxed selection on these genes (PBS regions - *O. ochracea* median log(*K*): 0.349; Background regions – *O. ochracea* median log(*K*): 0.244; *Mann-Whitney U* test, *p* = 0.46) (**Fig. 5F**). In fact, no species showed significantly relaxed selection on PBS genes, and in the majority the trend was in the other direction. Thus, genes involved in recent counter-adaptation in Hawaiian *O. ochracea* have been subject to episodic positive selection over the course of millions of years in related Tachinid fly lineages.

## Discussion

Novel interactions between species have the potential to escalate into coevolutionary arms races. However, the evolutionary dynamics and genetic signatures of these interactions in natural populations are obscure because detecting such events in ‘real-time’ is exceedingly difficult. Previous studies have shown that *T. oceanicus* populations exposed to a recently introduced endoparasitoid fly, *O. ochracea*, evolved a suite of signal-attenuating adaptations that protect against parasitism (*20*). Our results pinpoint the recent timing of this new species interaction using population genomic data, and reveal the genomic features of rapid counteradaptation in *O. ochracea*. The first adaptive host phenotype, ‘flatwing’, was detected in 2003, followed by at least four other adaptive wing morphs discovered as recently as 2019 (*20*, *23–25*, *35*, *58*). Genomic regions shaped by host-driven counterselection indicate that fly populations have been coevolving in lockstep with their hosts, potentially beginning ∼20 years ago when the first adaptive host morph emerged. Intense natural selection can drive reciprocal adaptation between newly introduced species, even after severe bottlenecks.

Our results extend the utility of classic theoretical models of coevolution—typically evaluated in canonical, extensively studied systems—to emerging coevolutionary interactions. We find evidence of directional positive selection in the form of genome-wide selective sweeps, consistent with an arms-race model of coevolution and an oligogenic basis of counteradaptation. This contrasts with many recent examples of antagonistic interactions that are better explained by balanced polymorphism and frequency-dependent selection at coevolving loci (*57*, *59–62*). Although arms-race dynamics have strong empirical support from genomic studies in plant–pathogen (*63–65*) and animal–virus systems (*66*, *67*), our study provides one of the few demonstrations of such dynamics in an animal–animal interaction, where candidate genomic loci associated with coevolving phenotypes in both species have been identified and the tempo of reciprocal adaptation is known (*30*, *32*, *34*). Whereas previous work on antagonistic coevolution has largely focused on traits linked to virulence and immunity (*8*, *66*, *67*), we show that traits involved in animal communication can experience similarly rapid, reciprocal evolution.

We combined gene-environment association analyses and selective sweep scans to identify two main candidate genes, *GRIK1* and *TLR3/Toll-7*, which partly underlie counter-adaptation to host-imposed selection in *O. ochracea* populations. Within *GRIK1*, an intronic allele associated with the intensity of host-selection has recently risen to high frequency (∼ 0.9) across Hawaiian populations and shows differential expression in neural tissues in gravid female flies upon hearing *T. oceanicus* song (*P*_adj_ < 0.001; logFC:-0.96), indicating a role for neural gene expression changes in *O. ochracea’s* counter-adaptation to increasing host silence. Structural or regulatory genomic changes underlying neural evolution and *O. ochracea’s* counter-adaptation in Hawaii is mechanistically plausible for a number of reasons. In *Drosophila*, *GRIK1* mediates motor control(*68*). Rapid evolution of it and similar proteins may have repurposed other sensory modalities to achieve acoustic host detection in *O. ochracea*. Ionotropic glutamate receptors may additionally function as chemoreceptors and rapid evolution of such proteins may imply repurposing other sensory modalities for host detection in *O. ochracea* (*69*, *70*). Similarly, whilst toll receptors are routinely involved in mounting immune responses, in flies *TLR3/Toll-7* are expressed in motor neurons and are required for normal locomotion (*56*).

It is intuitive to link this genomic result to the known heritability of neural responses to novel host songs (*37*), because covariation in fly responses and host song features has been demonstrated throughout *O. ochracea’s* range (*21*, *71*) and in other *Ormia* species (*13*). Our field-based gene expression assay showed that genes differentially expressed in neuronal fly tissues after hearing *T. oceanicus* song are also undergoing strong selective sweeps in Hawaii than would be expected by chance alone. Additionally, other selective sweep regions contain multiple genes with similar neural functions, for example a glutamate receptor *GRIN2B* (*Nmdar2*) linked to olfaction, synaptic function and plasticity (*50*), and *APP* (*Appl*) expressed primarily in *Drosophila* nervous systems and linked to multiple behavioural phenotypes (*48*, *49*). These results indicate that adaptive evolution of neural genes may be improving *O. ochracea’s* ability to detect novel morphs of *T. oceanicus,* or alternatively to diversify the hosts they target in Hawaii.

The demographic conditions that shape co-evolution influence both the likelihood and the speed of adaptation (*72*). For introduced species, establishment success and the ability to respond to biotic or abiotic selective pressures have been predicted to depend on levels of pre-existing genetic variation (*73–75*). Our results challenge this broadly accepted view. First, our genetic reconstructions validate anecdotal observations suggesting that *O. ochracea* was introduced to Hawaii within the last half-century (40). It has low effective population sizes, and consequently exhibits sharply reduced genetic variation relative to fly populations on the United States mainland. Similarly, analyses of *T. oceanicus* populations across Hawaii reveals congruent demographic patterns (*34*). Thus, reciprocal adaptation driven by escalating arms-race dynamics can occur in natural systems where neither host nor parasite are demographically stable, and where genetic variation is severely depleted. Our findings are in contrast to seminal experimental coevolution studies that use unicellular organisms with large effective population sizes, where adaptation is expected to proceed rapidly (*76*, *77*).

In coevolutionary interactions, it remains unclear whether coevolving loci driving contemporary adaptation have historically been under strong selection, perhaps because of long-term roles in mediating biotic interactions. Evidence from systems characterised by Red Queen dynamics shows that genomic signatures of balancing selection can persist for extraordinarily long periods (*57*, *78*, *79*), even between strongly reproductively isolated lineages (*80*). Signals of ancient or recurrent selective sweeps are more challenging to analyse over longer evolutionary timescales because recombination erodes regions of reduced genetic diversity (*8*). Taking advantage of recent analytical developments, we found that genes located within selective sweep regions in Hawaiian *O. ochracea* have repeatedly experienced episodic directional selection over the course of millions of years. Similar findings from animal–animal (*81*) and plant–herbivore (*82*) systems suggest that ancient sequence evolution at loci commonly involved in toxin resistance may enable rapid evolution on contemporary timescales, including between novel interactants. This historical contingency appears to play a similarly important role in bacteria–phage systems (*83*). The rapid coevolution of *O. ochracea* genes involved in counteradaptation to cricket defenses involved repeated co-option of the same genes for host detection across other tachinids, consistent with the remarkable diversity in tachinid sensory systems and host-detection strategies (*15*, *16*). Overall, our results provide insight into the genomic basis of rapid counter-adaptation and the turbulent evolutionary dynamics of an incipient coevolutionary interaction in the wild.

## Methods

### Sample collection, extraction and genomic sequencing

Whole-individual flies were collected from sampling locations across four islands across the Hawaiian archipelago (Kaua’i, Oah’u, Moloka’i and Hawai’i) using sound traps playing *T. oceanicus* song. Sound traps were constructed using MP3 players (SanDisk 368 SDMX26-008G-E46P) and Jam HX-P303BK speakers, playing a *T. oceanicus* song model constructed by averaging song parameters across several Hawaiian populations (*84*), at ∼85 dB at a distance of ∼5cm from the speakers. Hawaiian samples were collected across three field excursions in June 2022, October 2022 and June 2024. *O. ochracea* samples from California and Arizona were collected from Arizona and California in August 2022 using sound traps playing male advertisement song(s) from a range of *Gryllus* (*G. integer, G. lineaticeps, G. staccato, G. longicercus)* common known host species in Arizona and California (*22*, *85*) Individuals were flash frozen within an hour sampling and stored in 100% ethanol at-20C until DNA extraction. Finally, we also obtained outgroup (*Ormia depleta*) samples from Florida that were kept in 100% ethanol at-20C until DNA extraction.

DNA was extracted at the Tree of Life laboratory, Wellcome Sanger Institute (WSI). DNA was quantified and normalised to 200 ng in 120 µL. DNA was sheared to ∼450 bp using a Covaris LE220, and post-sheared samples were purified using SPRIselect beads. Libraries were constructed by performing end repair, A-tailing, and ligation using the NEB Ultra II custom kit. PCR was set up with KapaHiFi Hot Start mix and IDT UDI 96 PCR (sets A–D) barcodes, and six standard PCR cycles were performed. Libraries were uniquely dual indexed to mitigate tag hopping. PCR products were purified using SPRIselect beads, and final libraries were quantified using the Biotium Accuclear Ultra high sensitivity dsDNA Quantitative kit. Libraries were then pooled, normalised, and loaded onto the Illumina sequencing platform (Illumina NovaSeq 6000 and Illumina NovaSeq X).

### Genome assembly of O. ochracea

10 individuals of *O. ochracea* from a lab population (originally collected from Lai’e, Oah’u) were flash frozen and kept in-80C until high-molecular weight DNA extractions. Samples were then transported to the Sanger Institute in dry ice for high-molecular weight DNA and RNA extraction.

A single *O. ochracea* sample was weighed and dissected on dry ice with tissue set aside for Hi-C sequencing. Tissue from the same individual was disrupted to a fine powder using a Covaris cryoPREP Automated Dry Pulveriser and High molecular weight (HMW) DNA was extracted using the Qiagen MagAttract HMW DNA extraction kit. HMW DNA was sheared into an average fragment size of 12–20 kb in a Megaruptor 3 system with speed setting 30. Sheared DNA was purified by solid-phase reversible immobilisation using AMPure PB beads with a 1.8X ratio of beads to sample to remove the shorter fragments and concentrate the DNA sample. The concentration of the sheared and purified DNA was assessed using a Nanodrop spectrophotometer and Qubit Fluorometer and Qubit dsDNA High Sensitivity Assay kit.

Pacific Biosciences HiFi circular consensus DNA sequencing libraries were constructed according to the manufacturers’ instructions. DNA sequencing was performed by the Scientific Operations core at the WSI on a Pacific Biosciences SEQUEL II (HiFi) instrument. Hi-C data were also generated from head tissue using another *O. ochracea* sample from the same laboratory population using the Arima2 kit and sequenced on the Illumina NovaSeq 6000 instrument.

Assembly was carried out with Hifiasm (Cheng et al., 2021) and haplotypic duplication was identified and removed with purge_dups (Guan et al., 2020). The assembly was then scaffolded with Hi-C data (Rao et al., 2014) using YaHS (Zhou et al., 2023). The assembly was checked for contamination and corrected as described previously (Howe et al., 2021). Manual curation was performed using HiGlass (Kerpedjiev et al., 2018) and Pretext (Harry, 2022). The mitochondrial genome was assembled using MitoHiFi (Uliano-Silva et al., 2022), which runs MitoFinder (Allio et al., 2020) or MITOS (Bernt et al., 2013) and uses these annotations to select the final mitochondrial contig and to ensure the general quality of the sequence.

A Hi-C map for the final assembly was produced using bwa-mem2 (Vasimuddin et al., 2019) in the Cooler file format (Abdennur & Mirny, 2020). To assess the assembly metrics, the k-mer completeness and QV consensus quality values were calculated in Merqury (Rhie et al., 2020). This work was done using Nextflow (Di Tommaso et al., 2017) DSL2 pipelines “sanger-tol/readmapping” (Surana et al., 2023a) and “sanger-tol/genomenote” (Surana et al., 2023b). The genome was analysed within the BlobToolKit environment (Challis et al., 2020) and BUSCO scores (Manni et al., 2021; Simão et al., 2015) were calculated. The BRAKER2 pipeline (Brůna et al., 2021) was used in the default protein mode to generate annotation for the *O. ochracea* assembly. The X chromosome was identified by homology with *Panzeria rudis* (GCA_956483635.1).

### RNA-seq experimental design and sequencing

To identify genes involved in detecting *T. oceanicus*, we collected 16 gravid females of *O. ochracea* were collected from Common Ground, Kaua’i using sound traps. After a period of silence where no song was placed, we split the 16 gravid females into two groups, one exposed to 10 minutes of *T. oceanicus* song and the other to 10 minutes of silence. Both groups were immediately flash frozen in dry ice and subsequently heads were dissected from bodies and stored in RNAlater in screw-cap 1.5ml Eppendorf tubes for long-term storage before extraction.

Dissected heads were transferred into a 1.5 mL Eppendorf with of 300 µL TRIzol and homogenised via a hand-held homogenizer (Cole-Parmer® Motorized Pestle Mixer) for 5 minutes and left to incubate at room temperature for 5 minutes. Following homogenisation, we extracted total RNA using TRIzol (Invitrogen) and Qiagen RNAeasy kit. All samples were stored in-80°C. mRNA libraries were prepared using PolyA selection and sequenced on an Illumina NovaSeq 6000 to produce paired 150bp reads. Library preparation and sequencing was performed at Novogene.

### Mapping, variant calling and variant filtration

After initial quality control, reads were trimmed for quality and adapter sequences using *fastp* with default parameters (*86*). Reads were then aligned to the *O. ochracea* genome assembly using bwa-mem2 with default parameters (*87*, *88*). Mapped reads were then sorted, and PCR duplicates were flagged and removed using *Sambamba* (*89*). Read groups were then added to sorted, deduplicated mapped reads were removed using picard (*90*). Variant calling was performed using *bcftools* with all samples collected (*91*). Variants were subsequently filtered using *bcftools* to retain calls with minimum quality of 30, minimum depth of 10; site missingness of 10%; biallelic SNPs. Additionally, variants overlapping repetitive sequences inferred using program *Red* were removed using the intersect functionality in bedtools (*92*, *93*). Since the X chromosome consisted of mostly repetitive sequences (69%) where the autosomes showed much lower levels of repeat content (17-21%), we removed it from further analysis to ensure our interpretations of selection genome-wide were conservative (**Supplementary Fig 10**). Additionally, *O. depleta* samples were only included in one analysis of genetic differentiation since all *O. depleta* samples showed high levels of missingness (>70%) genome-wide. For this specific analysis, we only retained SNPs across all samples that were entirely non-missing (0% missingness) and conducted F_ST_ scans using 50kb windows and only analysed windows with a minimum of 10 SNPs.

### Population structure and demographic inference

To infer population structure and demographic history, variants were linkage-pruned using *PLINK* (parameters: indep-pairwise 50, 10, 0.1, v.1.9)(*94*). Principal component analysis of all *O. ochracea* samples from across sampling years was performed using *PLINK* and visualised using ggplot2(*95*). After observing strong clustering by year of Hawaiian *O. ochracea* samples, we explored phylogenetic relationships of using samples collected only in 2022, since the largest number of collections of mainland US and Hawaiian samples occurred from June – October 2022. We used a GTR model to infer a maximum-likelihood tree of *O. ochracea* samples using *FastTree* and used the R packages *ape* and *ggtree* to plot the inferred tree (*96–98*).

To further understand the demographic history of *O. ochracea* of Hawaiian and mainland US samples, we took three approaches. First, using the pruned dataset described above, we (1) inferred divergence history and split times between US mainland samples and Hawaiian samples using SFS-based inference approach *moments* within GADMA2 (*99*, *100*). We projected the site-frequency spectra down to 30 individuals from Hawaii and Mainland populations so as to prevent differences in sample size affecting our inference. Otherwise, we used default settings in *GADMA2* to perform demographic inference. Split times in genetic units were scaled to interpretable time estimates using a generation time of 1 month based on laboratory-based observations of generation time for *O. ochracea* (*39*), and a *Drosophila melanogast*er mutation rate of 3.32 x 10^-9^ (*101*).

Using the same pruned dataset, we (2) explored effective population size dynamics through time using *PHLASH*, a Bayesian implementation of PSMC approaches (*102*). Specifically, we inferred the demographic history of Hawaiian samples, Californian samples and Arizonan samples separately using the same linkage-pruned dataset as above, and visualised them using matplotlib. Finally, we (3) sought to understand the recent demographic history of Hawaiian *O. ochracea* samples using *GONE2*, which uses genome-wide patterns of linkage disequilibrium to reconstruct changes in N_e_ over the last ∼150 generations (*103*). Since we observed relatively weak population structure within the three Hawaiian Islands with the densest sampling (Kaua’i, O’ahu and island of Hawai’i), we grouped individuals from these islands together and performed island-level demographic reconstructions. Because we only collected five samples from Moloka’i we chose not to include these in recent demographic history reconstructions. We assumed a fixed rate of *ρ* = 1.5 cM/Mb, similar to the average recombination in *D. melanogaster* after accounting for the fact that there is no recombination in males (*101*). Recent demographic reconstructions using *GONE2* (and other linkage-based approaches) are dependent on recombination rate, so to assess whether our main conclusions for this analysis changed depending on recombination rate, we repeated the inference using slightly higher (*ρ* = 1.9 cM/Mb) and lower (*ρ* = 1.1 cM/Mb) fixed recombination rates and found this little affected N_e_ estimates. Separately, we also inferred recent demographic histories for individuals collected from Arizona and California using the same approach as a comparison point.

### Calculation of summary statistics and selection scans

#### Broad-scale descriptive statistics

To understand the consequences of demographic processes on genetic variation in Hawaii we calculated a number of summary statistics. Using VCFtools, we computed the inbreeding coefficient using filtered autosomal SNPs for each individual (*104*). To supplement inbreeding coefficients, we also calculated runs of homozygosity (ROH) and the fraction of the genome in runs of homozygosity (FROH) per island/sampling location using PLINK and R package detectRUNS (*105*). Additionally, using VCFtools we calculated Tajima’s *D* in 10kb in windows for each island in Hawaii and for US mainland populations separately, removing any windows which contained fewer than 100 SNPs. Similarly, using the same autosomal SNP dataset alongside invariant sites, we calculated π using pixy for Hawaiian samples and US mainland samples(*106*).

#### Selective sweep summary statistics/scans

To identify directional selection in Hawaiian populations of O. ochracea, we calculated the normalized population branch statistic (PBSn1). The PBSn1 statistic outperforms traditional F_ST_ scans and the population branch statistic at detecting local selective sweeps in simulations, particularly in the presence of background selection(*107*). We calculated three-way weighted F_ST_ between Hawaii, California and Arizona as the outgroup population to get pairwise distance metric (*Τ*) (Cavalli-Sforza 1969) (example below):

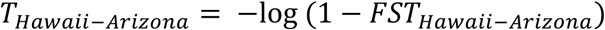

We then computed PBSn1 using the following equation:

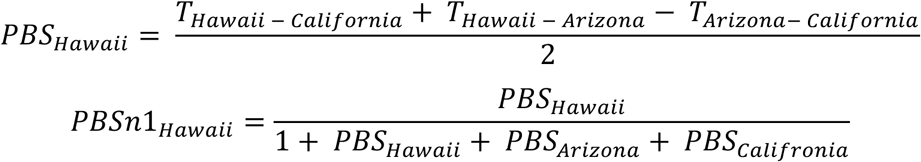

Alongside the PBSn1 statistic, we computationally phased our samples using *SHAPEIT4* using default settings and calculated XP-EHH to test for selection on Hawaiian versus US mainland samples using the R package *rehh* (*108*, *109*). Finally, we used SweeD, a maximum-likelihood implementation of CLR test, to detect selective sweeps on Hawaiian samples (*44*). For each test, we determined outliers using 95% percentile cut-offs. We identified conservative, candidate outliers for selective sweeps by intersecting outliers from all three of these approaches to detect selection. We performed functional enrichment for outlier genes using g:Profiler(*110*). Across all scans, we used individuals collected in the same field season to control for confounding effects of temporal genetic variation.

In the face of founder effects, genetic drift is expected to produce patterns of depleted genetic diversity that can strongly resemble directional selection captured in selective sweeps. To determine whether the specific demographic conditions in Hawaiian *O. ochracea* could explain patterns consistent with selective sweeps, we simulated a neutrally evolving population undergoing a severe bottleneck using SLiM (v.4.3)(*51*). Specifically, we simulated 10 Mb region with a uniform mutation rate of 1 x 10^-8^ and a recombination rate 1 x 10^-6^ and only neutral mutations. Additionally, we started the simulation of with a population size of 100,000, where for 9,400 generations the population size stayed constant and at generation 9,400, the population experienced a severe bottleneck reducing it to only 5,000 individuals, which remained unchanged until the end of the simulation at 10,000 generations. This simulation closely mirrors the demographic scenario identified in the observed data for Hawaiian *O. ochracea*. We outputted variants for 186 individuals into a VCF file, representing the same number of Hawaiian *O. ochracea* individuals identified in our 2022 sampling season and in our final, filtered dataset. We re-ran SweeD on this simulated data using default settings.

We used SINGER (*111*) to perform ancestral recombination graph (ARG) reconstruction using phased *O. ochracea* samples on chromosome 2 specifically, where candidate genes under directional selection are located. Specifically, ARG inference was performed using *SINGER* using the following parameters “-m 2.8e-9-Ne 1e5-n 100-thin 20” to describe an approximate mutation rate for *O. ochracea* and an effective population size of 100,000 which spans the effective size of *O. ochracea* on the mainland and in Hawaii. These values represent a best guess at the mutation rate and effective population size, but *SINGER* accounts for model misspecification and unmodeled changes in effective population size. Additionally, our choice of *SINGER* for ARG reconstruction was two-fold: (a) *SINGER* produces remarkably accurate ARG inferences compared to other state-of-the-art tools (*Relate, tsinfer ARGWEAVER*) and (b) is most efficient with hundreds of samples rather than tens of samples or thousands of samples(*111*).

We inferred selection coefficients and allele frequency trajectories for SNPs identified in gene-by-environment analyses that overlap with selective sweep scans. These SNPs are associated with three genes (*GRIK1*, *TLR3* and an unannotated gene). We focus on the two candidate SNPs within the two annotated candidate genes and infer selection coefficients for them. Inference using CLUES2 (*112*, *113*) was performed using the “inference.py” functionality with the following parameters “--N 100000 --tCutoff 10000 --CI 0.95” to specify the approximate effective population size (consistent with the SINGER inference), the maximum generation cut-off time and specifying confidence 95% intervals. We performed this analyses for three time-epoch models which allows for estimation of selection coefficients across a user-defined time period. We estimated the strength of selection using a one-epoch model where selection pressure (*s*^MLE^) does not vary from current day through to 10,000 generations ago, a two-epoch model where *s^MLE^* varies between the first specified epoch (1,000 – 10,000 generations ago) and the second (0 – 1,000 generations ago), and finally, a three-epoch model where *s^MLE^*varies between the first (1,000 – 10,000 generations ago), second (200 – 1,000 generations ago) and third-epoch (0 – 200 generations ago). We include parameter estimates and likelihoods for all models in **Supplementary Table 4**. Additionally, for these two SNPs we inferred the historical allele frequency trajectory using CLUES2 and the “plot_traj.py” script. This allowed us to determine approximately when allele frequencies in both variants changed through time.

#### McDonald-Kreitman tests and Direction of Selection

To further investigate the evidence for positive selection in genes within candidate selective sweeps, we asked whether we could detect higher levels of adaptive evolution in this subset of genes versus the rest of the genome. To do this, we downloaded short-read data for *Tachina fera*, a parasitoid fly within the same sub-family as *O. ochracea*. We aligned *T. fera* short-read data to *O. ochracea’s* genome using the same genomic mapping pipeline as above, and variants called using the exact same protocol as above via *bcftools*. We then merged the variant data containing the *T. fera* sample to our filtered VCF containing samples of *O. ochracea* from Hawaii and North America. We then computed gene-wide polymorphism (*pN* and *pS*) and divergence (*dN* and *dS*), the direction of selection (DoS) statistic and performed McDonald-Kreitman tests using the software package *degenotate* (*114*). Specifically, we computed these statistics and performed tests separately for Hawaiian and North American populations of *O. ochracea*, so that patterns of adaptive evolution could be contrasted between the two.

#### Gene-by-environment association test

Alongside *O. ochracea* sampling, we performed *T. oceanicus* male morph surveys in 12 focal populations across Hawaii. At each population, we searched and collected crickets using a team of two to three persons for 1-2 hours, whilst simultaneously collecting *O. ochracea* using playbacks. After collection, we counted the number of individuals collected and determined sex and morph abundances (based on sampling expertise and photographic comparisons). At all sites where we collected individuals that appeared to be singing-capable, field observations were used to verify that songs were heard on-site. To account for inherent unsampled variation, instead of using quantitative per site measures of singing-capable males, we instead categorised sites into three specific groups using the morph abundance data: sites with high-intermediate representation of singing-capable individuals (designated 1.0), sites with intermediate-low representation of singing-capable individuals (designated 0.5) and sites with no singing-capable individuals (designated 0.0). Sites where we noted singing-capable males but could not sample any because of cricket avoidance behaviour (i.e. burrowing) were included in the intermediate-low representation category.

We tested whether regions with genomic signatures consistent with selective sweeps were related to loss of song in *T. oceanicus* populations using latent factor mixed models (*LFMM2*) and our categorical measure of singing-capable males described above as a response variable (*115*). This approach tests for SNPs differing in allele frequency that are associated with the proportion of singing males whilst accounting for hierarchical population structure across populations and islands. SNPs were pruned using PLINK (*--indep-pairwise 50 10 0.1*) to ensure each SNP could be considered as independent from each other. Missing genotypes were imputed using the *strimpute* function in LEA and to estimate population structure, where K specified as K = 3 following examination of elbow plots. Parameter estimation in LFMM was performed using ridge regression(*116*). SNPs were considered significant if the adjusted (Bonferroni-Hochberg) p-value < 0.05.

#### Differential expression analysis

Raw paired-end RNA-sequencing reads were collected from heads of *O. ochracea* exposed to silence and *T. oceanicus* song. Raw reads were trimmed and filtered using *fastp* with default parameters and then aligned to the *O. ochracea* genome using STAR (*117*), with the following parameters: “--genomeSAindexNbases 13”; “--sjdbGTFfile $gff” and “--sjdbOverhang 149”. Across samples, unique mapping rates ranged from 87-91%. Gene counts were estimated from BAM outputs from STAR using RSEM (*118*), with the following paraemeters: “--estimate-rspd”; “--append-names”; “--seed 123456” and “--no-bam-output”. We imported gene counts and quantified differential expression between individuals exposed to silence and *T. oceanicus* song (∼ condition) using DESeq2 (*119*). To account for hidden variation due to biological and technical confounds, we visualised expression of samples using PCA and conducted surrogate analysis using svaseq (*120*). We adjusted our model to include surrogate variables explaining hidden variation (∼ condition + SV1 + SV2 + SV3 + SV4). Additionally, we applied shrinkage of the log_2_ fold change estimates to obtain more accurate fold change estimates using the package *apeglm* (*121*). Differential expressed genes were identifying using FDR-adjusted p-values at a cut-off of 5%

### Comparative genomics of Tachinidae

Nine chromosome-level genomes of other tachinids used in this analysis were produced by the Darwin Tree of Life project. We retrieved these genomes from NCBI, where genomes from the Tree of Life project are deposited. To assess the quality of genomes, the relationship between different species within the family Tachinidae and their chromosomal evolution, by identifying conserved single-copy orthologs via Benchmarking Universal Single-Copy Orthologs (BUSCO) (*122*, *123*). Genome quality was assessed using BUSCO (v5), specifying the Diptera (diptera_odb10) reference gene set (n = 3,285) for each of the nine tachinid species. We assessed the proportion of single copy orthologs recovered for each species to identify species with high levels of duplication or missingness, and analysed metadata on genome and chromosome sizes across the additional nine species. A list of the species used in the analysis and genome statistics can be found in **Supplementary Table 5** and shown in **Supplementary Fig 6-9**.

To infer species relationships, we used species-specific proteomes and *Orthofinder* to recover 1,452 single-copy orthologs that were present in all species and were used for species tree estimation(*124*). Each of the 1,452 single-copy orthogroups were aligned using *MAFFT* using the default parameters and gene trees for each aligned, single-copy orthogroup were generated using *FastTree* (*125*, *126*). We used the *STAG* approach within Orthofinder to infer the species tree. Instead of relying only on single copy orthologs, *STAG* uses multi-copy orthogroups to infer species trees and shows comparable performance to other species tree inference approaches (e.g. concatenation and multi-species coalescent inference) (*127*).

For molecular evolution analyses, we repeated our *Orthofinder* approach but with available coding sequences from each species of the nine tachinid species as well as *O. ochracea.* Using this approach, we identified 5,997 number of family-level orthologous alignments, which we focused on for further molecular evolutionary analyses. We then converted coding sequence alignments to protein alignments using *MACSE* (v2) and the “translateNT2AA” functionality, and protein alignments for all orthogroups were then aligned using *MAFFT* with default parameters(*128*). We then used *PAL2NAL* to convert the aligned protein sequence back into codon alignment disallowing gaps or mismatching sequences between the coding sequence and protein alignment using the parameter “-nomismatch” (*129*). These codon alignments were used for all *dN/dS* based analysis via the HYPHY framework. Alongside codon alignments, we used orthogroup-specific gene trees generated by *Orthofinder*, via *FastTree* as above, for all branch-based tests of molecular evolution.

We employed three tests within the HYPHY hypothesis-testing framework to assess lineage-specific positive selection using the *dN/dS* metric. These were: (1) *BUSTED-PH*, which tests for positive selection at one or more codons in specified ‘foreground’ and ‘background’ branches (*130*, *131*); (2) *RELAX*, which evaluates whether selection has been relaxed or intensified on foreground branches relative to background branches(*132*); and (3) *FUBAR*, which detects site-wise signatures of positive selection across a phylogeny(*133*). We tested whether the 348 genes identified as outliers in our normalised population branch statistic (PBSn1) analysis showed evidence of positive selection across tachinids. For *BUSTED-PH* and *RELAX*, the foreground branch was defined as the terminal branch leading to *O. ochracea*, with all remaining branches treated as background. For *FUBAR*, positive selection was inferred when at least one site showed a posterior probability > 0.9.

Our main aim was to determine whether these locally adaptive genes exhibit positive selection uniquely in *O. ochracea*, or whether similar selective pressures act across tachinids more broadly. As a result, we first examined *BUSTED-PH* results: this method tests for episodic positive selection on foreground and background branches separately and then compares their ω (*dN/dS*) values using a likelihood-ratio test to determine whether the two branch partitions differ significantly. We next used *FUBAR* to confirm whether genes identified as positively selected by *BUSTED-PH* also contained at least one site under positive selection. Finally, for genes showing significant differences in ω between branch partitions, we applied *RELAX* to assess whether these patterns were driven by relaxed selection (k < 1) or intensified selection (k > 1) on the branch of interest.

## Acknowledgements

We are very grateful to all landowners at all locations across Hawaii for permission to collect crickets and parasitoid flies on their grounds. We thank the Wellcome Sanger Institute Tree of Life Core Laboratory and the Long Read Extraction team at Sanger for DNA extraction and preparation for sequencing, Sanger Scientific Operations for long read sequencing and Illumina sequencing, and the Tree of Life Assembly team for primary genome assembly.

## Author contributions

Conceptualization: LHY, MB, XZ, SB, RMT, DAG, NWB

Methodology: LHY, JGR, RZ, MB, DAG, NWB

Investigation: LHY, JGR, RZ, KZT, SB, RMT, DAG, MP, NL, NWB

Visualization: LHY

Funding acquisition: XZ, MB, NWB

Project administration: NWB, LHY

Supervision: NWB

Writing – original draft: LHY

Writing – review & editing: LHY, JGR, RZ, KZT, NL, RMT, DAG, MB, NWB

## Data and materials availability

*O. ochracea* genome is available on NCBI under the BioProject accession PRJEB65437. Genome annotation for O. ochracea can be found on XX using the following link: https://ftp.ensembl.org/pub/rapid-release/species/Ormia_ochracea. Whole genome resequencing of O. ochracea is available on NCBI SRA under accession PRJEB63617. RNA-seq data can also be found on NCBI SRA under the accession PRJNA1422981. Scripts for population genomic analysis can be found on GitHub: https://github.com/LeebanY/EcologicalGenomicsOfNovelHostParasiteInteraction_Ormia_ochracea_MS.

