## Supplementary Text/Material for "Ecological genomics of a novel host-parasitoid arms-race in nature"

S1. Protein-coding genes under local selection in Hawaii show stronger evidence of adaptive evolution.

We further explored the evidence of strong directional selection in protein-coding genes in selective sweeps specific to Hawaiian *O. ochracea* populations by contrasting levels of polymorphism and substitutions to test for adaptive evolution. We leveraged chromosome-level genome and whole-genome resequencing data of the species *Tachina fera*, which belongs to the same sub-family (*Tachininae*) as *O. ochracea,* to calculate the proportion of adaptive substitutions ($\alpha$) and direction of selection (*DoS*) in genes implicated in candidate sweep regions. Positive values of *DoS* imply adaptive evolution of protein-coding genes, whilst negative values suggest purifying selection. Given that we have identified a cohort of genes involved in both adaptation and speciation in *O. ochracea*, presumably driven by co-evolutionary interactions, we predicted that this cohort of genes may be repeatedly under selection across macroevolutionary time. Additionally, levels of adaptive evolution are only expected to differ between Hawaiian and Mainland populations of *O. ochracea* due to contemporary changes in selection given the recency of divergence between these populations.

Overall, *DoS* is negative across all genes surveyed (n=11,603) implying a preponderance of slightly deleterious mutations and that genes undergoing adaptive evolution in both Mainland and Hawaiian *O. ochracea* populations are enriched for RNA metabolism ($p$ = 0.009) and cell cycle ($p$ = 0.01) processes (**Supplementary Fig. 3**). We find that *DoS* for genes in selective sweep regions in Hawaiian populations is significantly higher than compared to the genomic background (two-tailed Welch's two-sample $t$-tests: $p = 0.034$), whilst the *DoS* in selective sweep regions in mainland populations is not significantly different from the background (two-tailed Welch's two-sample $t$-tests: $p = 0.074$). We find stronger patterns of adaptive evolution in protein-coding genes within (5%) and outside (9%) of sweep regions in Hawaiian populations relative to mainland populations (sweep regions: 4%; background: 5%) of *O. ochracea* (**Supplementary Fig. 3**). Consistent with our selection scans, our results suggest higher rates of adaptive evolution in protein-coding genes in Hawaiian populations in genes within sweep regions, and across the genome, compared to mainland populations.

S2. Little evidence for balancing selection in parasitoid fly populations.

Hawaiian populations of *O. ochracea* show signals of strong directional selection, with little evidence of balancing selection, providing evidence for arms-race coevolution over Red Queen dynamics. However, contrasting genomic signatures of balancing selection versus directional selection in Hawaiian populations of *O. ochracea* is hampered by the fact that pronounced population contractions produce intermediate haplotypes genome-wide, confounding genomic tests of balancing selection (**Fig 2C**). North American populations of *O. ochracea* show little evidence of population structure or demographic fluctuations that have affected genetic diversity (**Fig 2B-G**). Using North American populations of *O. ochracea*, we asked if regions linked to host detection in Hawaiian *O. ochracea* populations show evidence of balancing selection in North American populations. Using genomic regions (n=67) identified across all three selective sweep statistics (i.e. SweeD, XP-EHH, *PBSn1*) in Hawaiian populations, we assessed whether the same genomic regions are (a) not under selection, (b) under directional selection or (c) under balancing selection in demographically stable North American populations. We found little evidence for increased Tajima's $D$ in genomic regions locally selected in Hawaii compared to the genomic background in Californian (two-sample t-test: t = -1.48, $df = 488$, $p = 0.13$) and Arizonan (two-sample t-test: t = -1.59, $df = 479$, $p = 0.11$) populations. In fact, we found that more of the 67 tested genomic regions showed depleted genetic variation ($D < -1$; 9.7% and 4.7% of genomic regions in Arizonan and Californian populations, respectively) compared to locally selected genomic regions with excess intermediate haplotypes characteristic of balancing selection ($D > 1$; 0% and 0.01% of genomic regions in Arizonan and Californian populations).

Similarly, we found no evidence for extreme nucleotide diversity ($\pi$) in these 67 genomic regions compared to the genomic background in Californian (two-sample t-test: t = 4.22, $df = 275.13$, $p < 0.001$; mean $\pi$ genomic background: 0.007, mean $\pi$ in locally selected genomic regions: 0.006) and Arizonan (two-sample t-test: t = 4.13, $df = 275.15$, $p < 0.001$; mean $\pi$ genomic background: 0.007, mean $\pi$ in locally selected genomic regions: 0.006) populations. North American populations show elevation in *D and* $\pi$ around centromeres and on the first ~30 Mb of the first autosome (**Supplementary Fig. 4**), potentially indicating a structural variant specific to North American populations. Overall, our results imply that Red Queen dynamics are unlikely to be driving adaptation and counter-adaptation in *O. ochracea*, consistent with theoretical models of host-parasitoid evolution.

### S3. Estimation of selection coefficient and historical allele frequency trajectories for alleles in candidate genes that are significantly associated with host selection.

Next, we sought to explore the evolutionary history of variants associated with rapid counter-adaptation in *O. ochracea* to increasing host silence. In particular, we estimated the selection coefficient and onset of selection in two variants (chr2: 58561647 and chr2: 58829212) associated with *GRIK1* and *TLR3*, our main candidates, by reconstructing the ancestral recombination graph at these loci and using a full-likelihood method, CLUES2, to infer the selection coefficients and onset of selection at variants associated with local adaptation. These two variants were selected because they fall within genic regions of *GRIK1* and *TLR3*, so we could be sure that selection coefficients could only be attributed to those two genes. Other intergenic SNPs close-by *TLR3* were of interest, but we could not be sure that they did not affect gene expression of another locus nearby or were not trans-acting.

We found exceptionally strong selection ($s^{MLE}$) for both variants (**Supplementary Table 4**) in a one epoch model, where a single selection coefficient was estimated from 10,000 generations to the current day ($s^{MLE} = 0.028 – 0.038$). Allele frequency trajectories for the derived allele at these two variant positions, and particularly for the intronic variant in *GRIK1* (chr2: 58561647), we observe a sharp increase in allele frequency around ~50 years ago (**Supplementary Fig. 5**) (generations scaled by a generation time of 12 generations a year and a mutation rate of $2.8 \times 10^{-9}$). These analyses provide further evidence for an incredibly strong and recent selective sweep linked to host adaptations.

### S4. Comparative genomics uncover genome dynamics across tachinids.

We originally downloaded all 18 tachinid genomes that were available and sequenced to chromosome-level by the Tree of Life programme but randomly subsampled the number of species used to reduce computational complexity of comparative analyses (particularly ortholog detection). In all analyses, but the synteny analyses, which included a subset of six of the tachinid genomes chosen to represent phylogenetic breadth based on the most recent tachinid phylogeny(*134*), the same ten randomly subsampled genomes were used. Across the ten subsampled genomes used (including *O. ochracea*), the genomes showed high levels of contiguity and completeness determined by recovery of single-copy orthologs present across dipteran genomes (**Supplementary Fig. 6**). Tachinid genomes are varied in genome sizes, but we find that this occurred via fairly equal reduction and/or expansion in genome content across chromosomes (**Supplementary Fig. 7**). Synteny analysis revealed that tachinid genomes show inter-chromosomal conservation, but considerable within-chromosome gene reorganization (**Supplementary Fig. 8**).

Evolution of repeat content appears to be fairly dynamic across *Tachinidae*. For example, we observed reductions in repeats across all autosomes in *O. ochracea,* potentially explaining the difference in genome size in *O. ochracea* compared to other tachinids (**Supplementary Fig. 9**). The middle of each autosome showed a pronounced increase in repetitive content, indicating metacentric centromeres are common in tachinids. Across tachinids, the X chromosome appears to be gene poor and characterised by high levels of repetitive sequence (**Supplementary Fig. 10**), consistent with genetic degeneration. For this reason, we did not consider the X chromosome in any population genomic analyses.

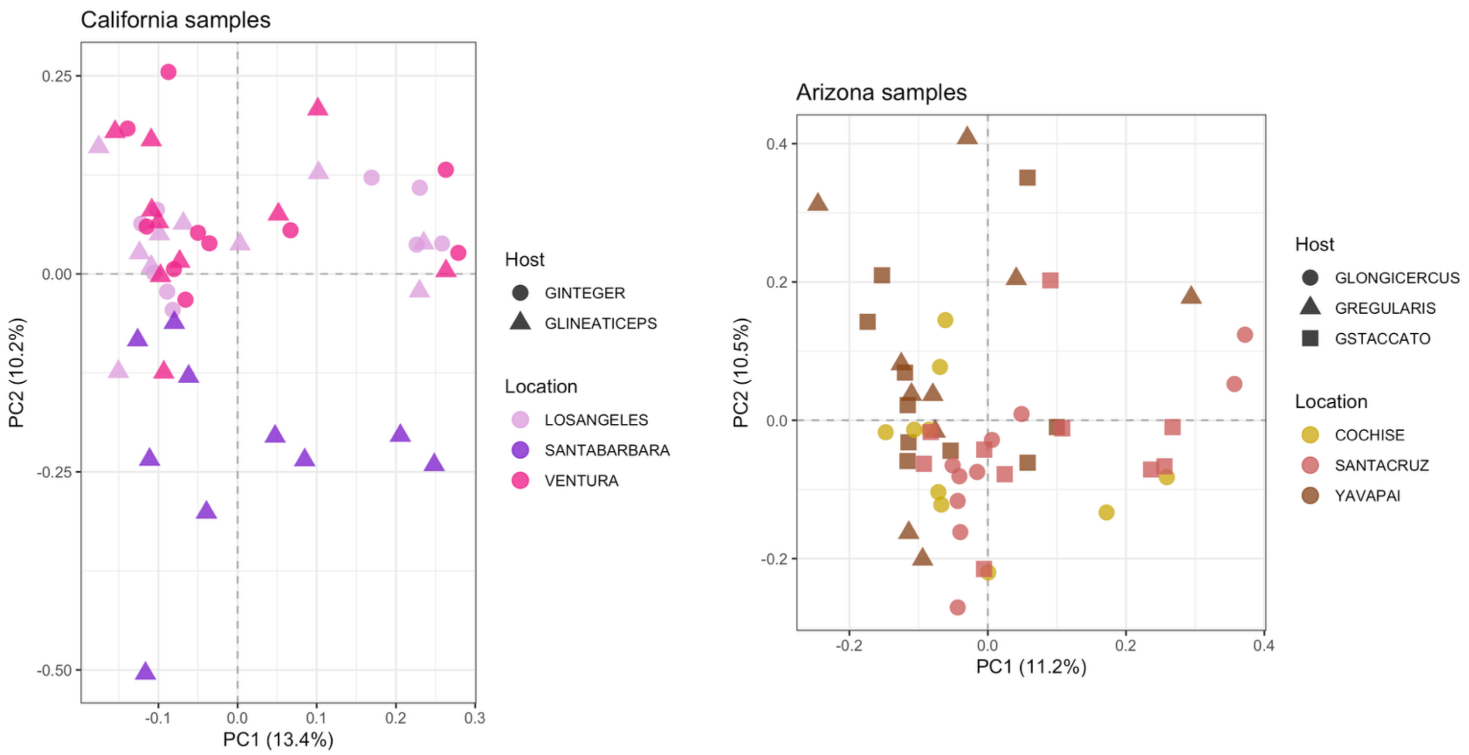


**Supplementary Figure 1:** PCA showing genetic variation sampled for *O. ochracea* samples collected from across California and Arizona in 2022 (n=99). Linkage-pruned autosomal SNPs used for PCA. Colours denote county/area where the samples were collected from whilst shapes denote the songs used during playbacks to catch each fly. PCA shows little evidence of cryptic differentiation of O. ochracea individuals by host species and weak evidence of population structure, matching patterns observed in Hawaii.

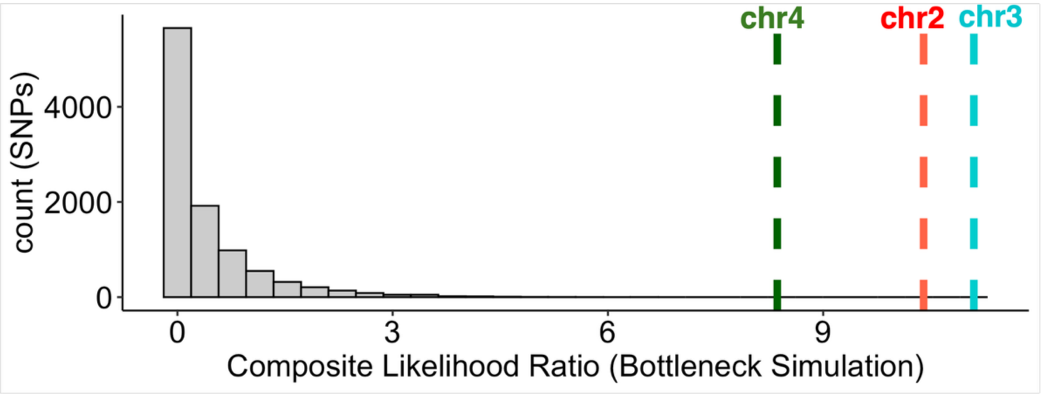


**Supplementary Figure 2**: Composite likelihood ratio for bottleneck simulation modelled based on the evolutionary history of Hawaiian *O. ochracea* populations. A 10 Mb genomic sequence consisting of neutral mutations was simulated with an initial population size of 100,000 that was significantly reduced to 10,000 individuals due to a recent, founder effect. Genetic variation for 186 simulated individuals was outputted and used to estimate the composite likelihood ratio test to test for a selective sweep at each SNP in the simulated dataset. The distribution reflects CLR values from this simulation. The coloured dashed lines are mean CLR values at observed outlier loci on chromosomes 2, 3 and 4. Demographic conditions of Hawaiian *O. ochracea* populations cannot, on their own, explain signatures of selective sweeps in the observed data.

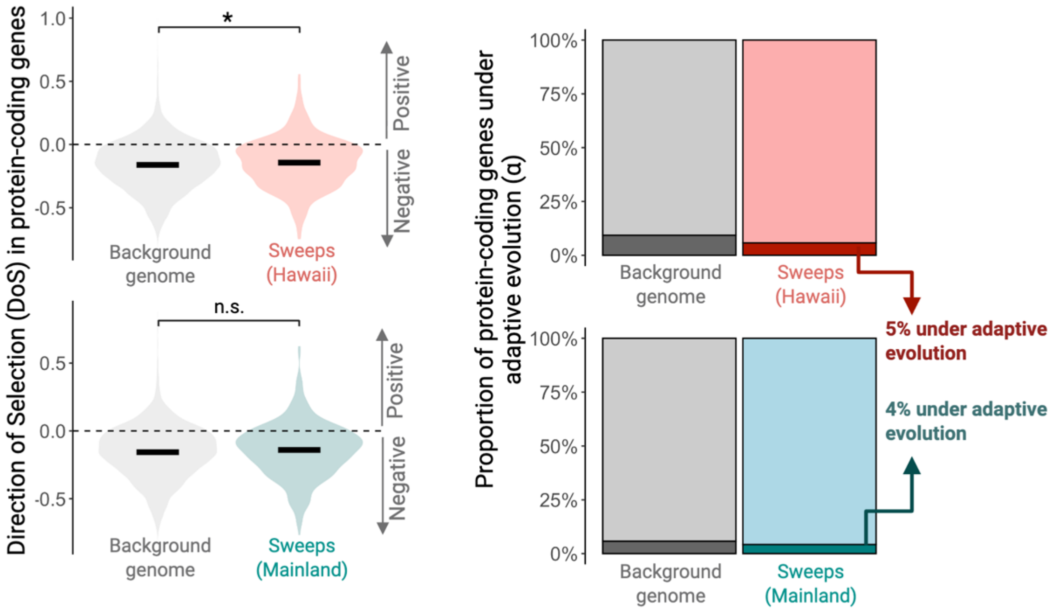


**Supplementary Figure 3**: Using an outgroup species of parasitoid fly (*T. fera*), we calculated the Direction of Selection (DoS) and performed McDonald-Kreitman tests for all protein-coding genes to determine support for long-term adaptive evolution of genes within selective sweeps in Hawaiian populations of *O. ochracea*. We compared the DoS for genes found in Hawaii-specific selective sweeps in Hawaiian populations (red violin plot) and in the mainland US populations (blue violin plot) and compared them to the genomic background (all other protein-coding genes). Negative DoS denotes negative/purifying selection, whilst positive values imply positive selection. Similarly, we calculated the proportion of protein-coding genes within Hawaii-specific sweeps with significant McDonald-Kreitman tests after multiple testing correction (FDR < 0.05) in Hawaiian populations (dark red) and in the US mainland (dark blue).

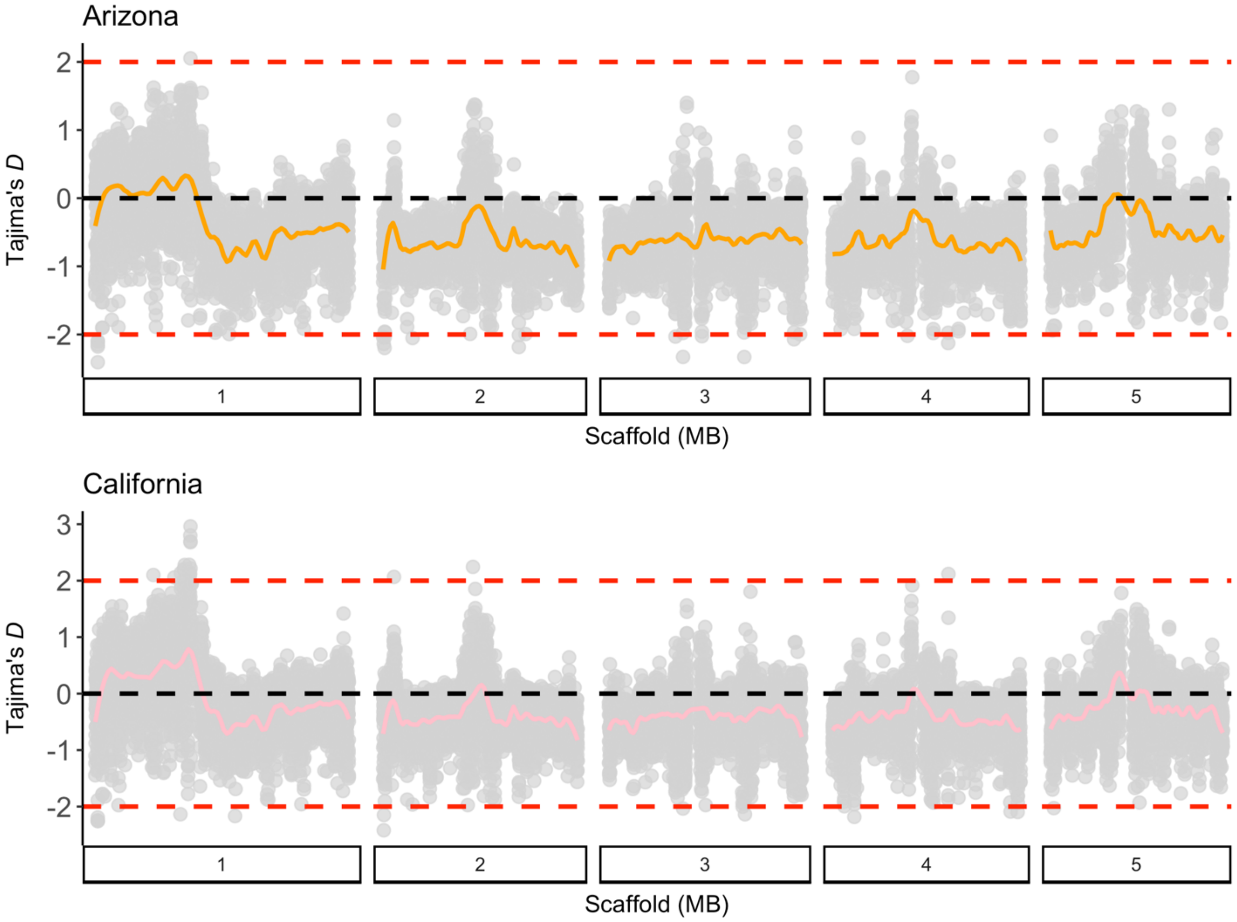


**Supplementary Figure 4**: Patterns (smoothed in orange and pink) of Tajima's *D* in Californian and Arizonan populations of *O. ochracea*. Regions of extreme *D* are denoted by the red dashed lines. Regions of increased genetic variation (positive *D*) are concentrated on the first half of the first autosome and in potential centromeric regions across the genome.

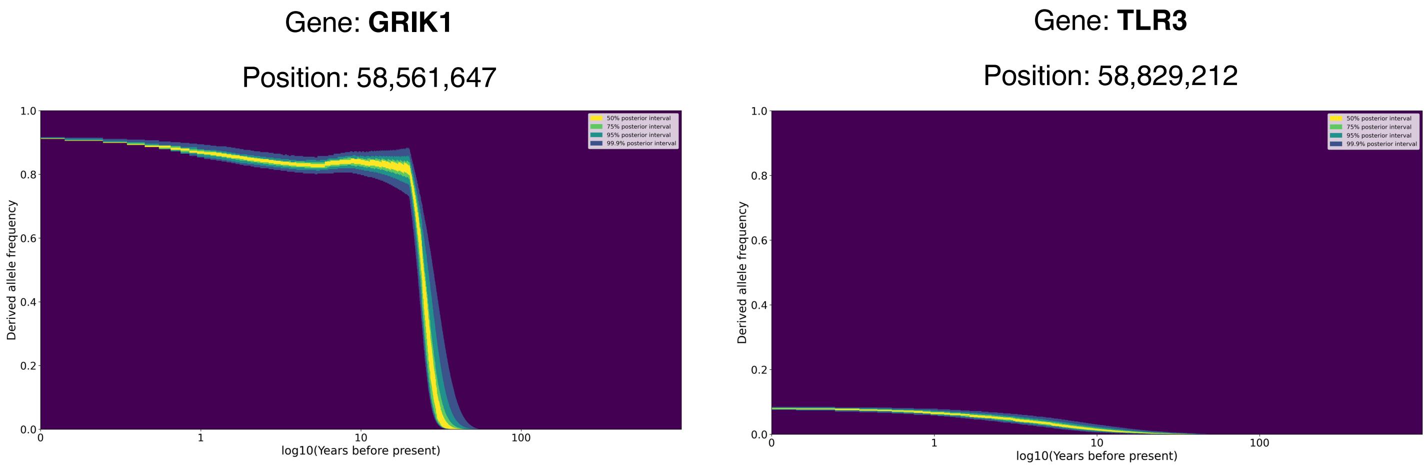


**Supplementary Figure 5**: Heatmaps showing the inferred posterior density of the derived allele frequency back in time (on a log$^{10}$ scale) at two candidate alleles found in two candidate genes (GRIK1 and TLR3), estimated using *CLUES2*.

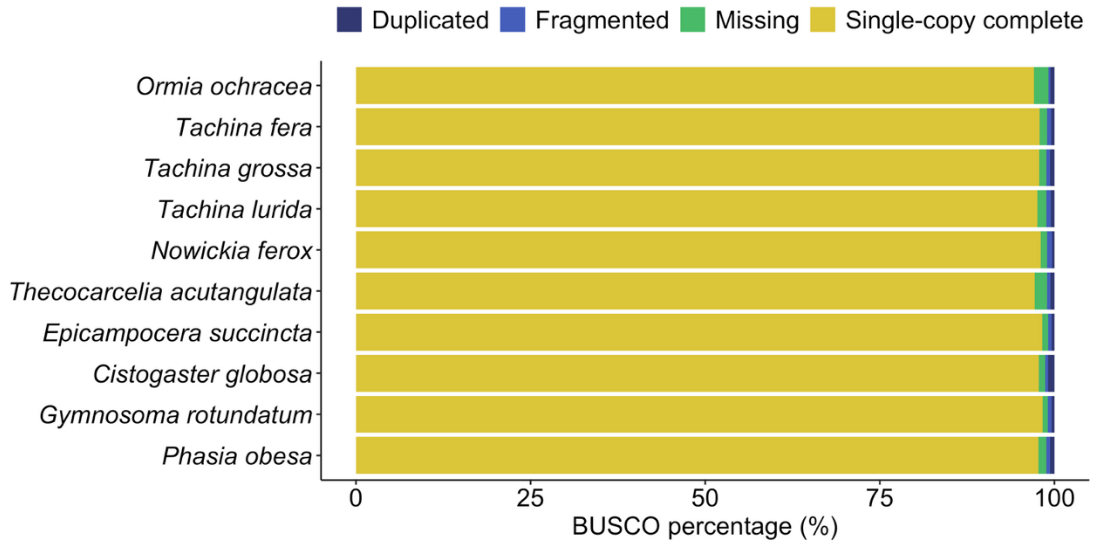


**Supplementary Figure 6**: Assessment of genome completeness for tachinids used in comparative genomic analyses. Tachinid genomes produced by the Tree of Life sequencing effort are incredibly contiguous, with most genomes showing almost complete recovery of single copy orthologs found in dipteran genomes.

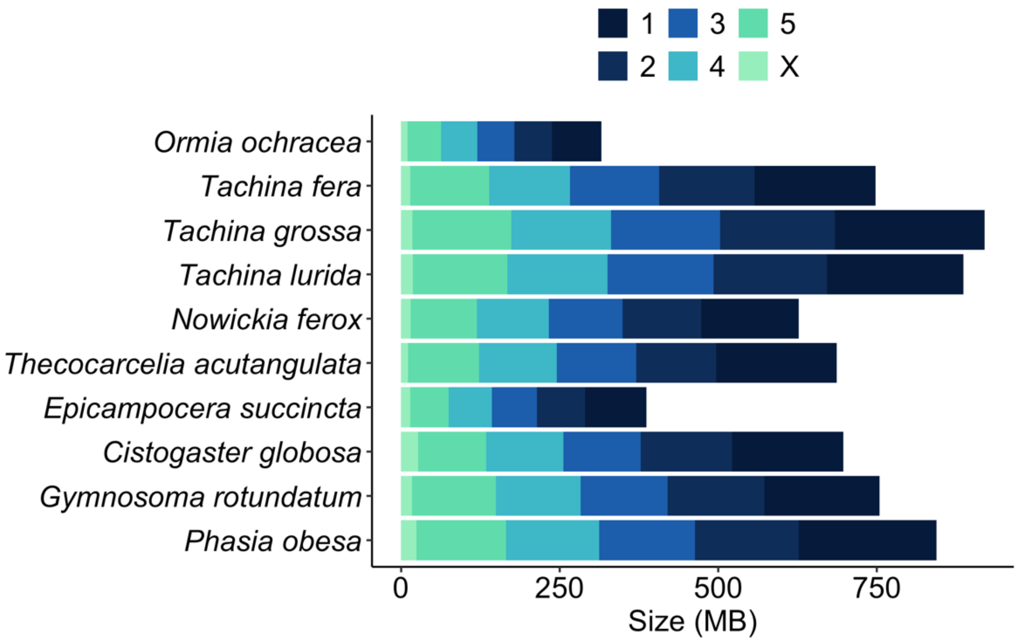


**Supplementary Figure 7**: Considerable variation in genome and chromosome sizes between tachinids considered in molecular evolutionary analysis. Reduction or expansions in genome sizes appear to occur as a result of changes in genomic content across all chromosomes, apart from the X chromosome, which is small and stable across tachinids.

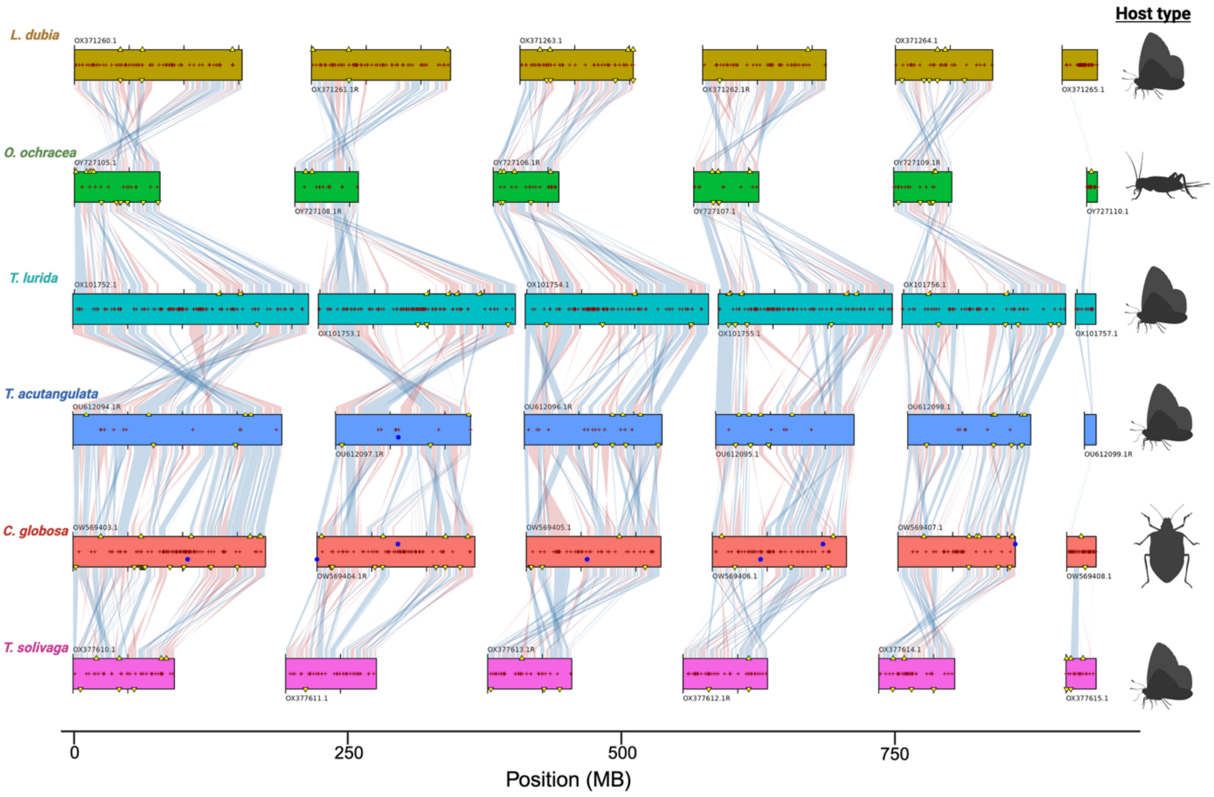


**Supplementary Figure 8**: Conserved synteny across tachinids inferred from dipteran BUSCO genes. Synteny blocks showing collinear "Complete" BUSCO genes connect scaffolds from adjacent assemblies: blue indicates genes on the same strand; red indicates genes on the opposite strand. Filled blue circles show predicted telomeres from the tool *TIDK*. Assembly gaps are indicated by dark red ± symbols. Duplicated BUSCOs are represented by yellow triangles. Genes on the forward strand are positioned above the center line, while reverse strand genes are below. Chromosome lengths are scaled relative to the total genome size of each assembly (scale = pc), normalizing for size differences and facilitating comparisons between genomes with markedly different chromosome sizes. Host types for each species are shown based on life-history information found in references in Supplementary Table 5.

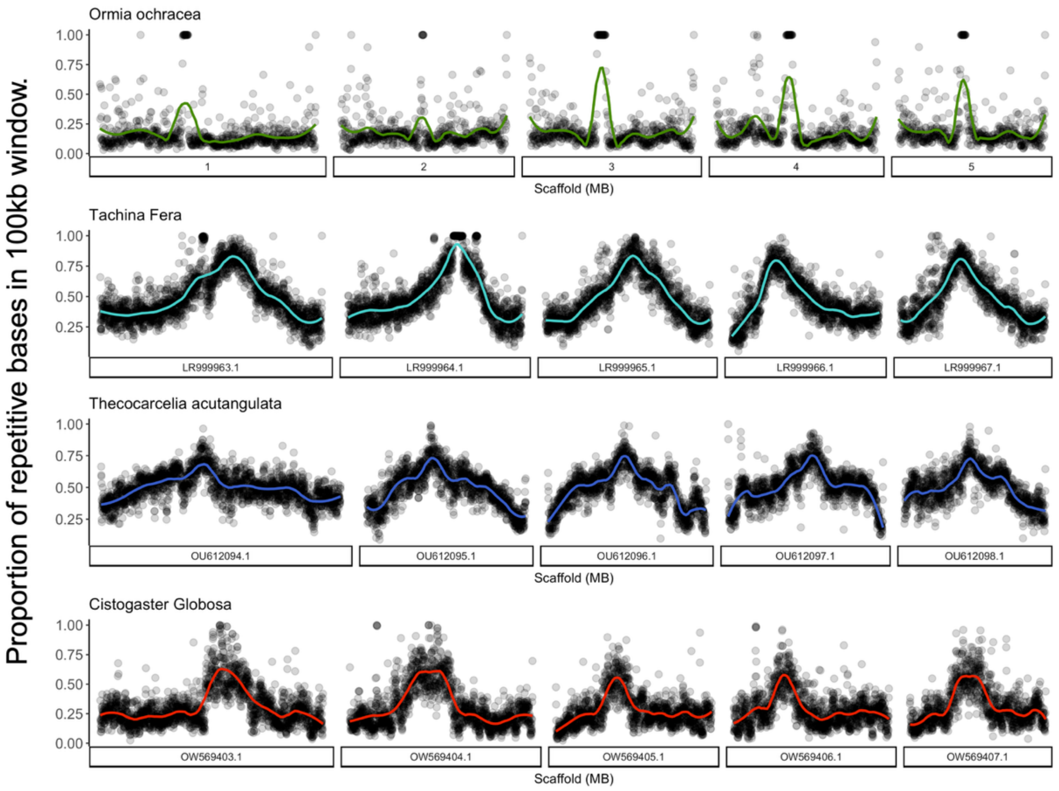


**Supplementary Figure 9**: Repeat landscape shows genome-wide reduction in repetitive bases in *O. ochracea* compared to other tachinids. Repeats were inferred using Red, which is able to detect transposons and simple repeats and then plotted by the proportion of repetitive bases in 100kb.

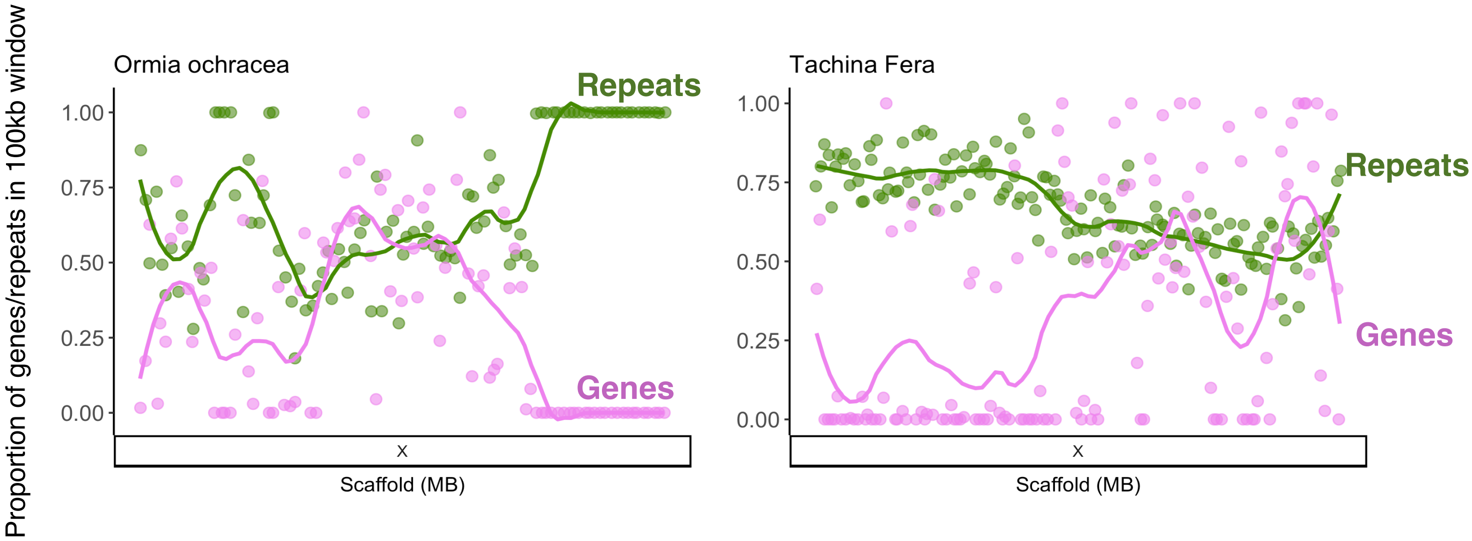


**Supplementary Figure 10**: Showing the proportion of repeats, inferred by *Red*, and genic regions in 100kb windows on the X chromosome for *O. ochracea* and *T. fera*. The X chromosome is highly repetitive at the beginning of X chromosome in *T. fera*, whilst the reverse is true in *O. ochracea*. Regions of high repetitive elements are characterised by remarkably few genes.

**Supplementary Table 1**: Field survey of *T. oceanicus* singing-capable and adaptive morphotypes across sites in Hawaii collected in 2022 and used for gene-by-environment analysis. Site labelling follows the convention of island and then abbreviated site names. 'Nw_males' correspond to the number of singing-capable males; 'CwNw' correspond to individuals with intact sound-producing structures but with curly wings; 'SwNw' correspond to individuals with intact sound-producing structures but small wings; 'Fw' corresponds to individuals with reduced or no sound-producing structures; 'CwFw' individuals have reduced or no-producing structures and curly wings. The number of *O. ochracea* individuals collected and sequenced from these sites at time of host surveys. We then binned sites into categories according to whether they contained no singing-capable males (0), low quantities of singing-capable males (0.5) or intermediate/high quantities of singing-capable males (1). Sites where 'Nw_male' contains an asterisk indicate locations where singing males were recorded in field notes but not collected. These sites were therefore classified as 0.5. These categories were used to test for an association between genomic regions and host-imposed selection on *O. ochracea* using a gene-by-environment association test.

| **Site** | **Nw_male** | **CwNw_male** | **SwNw_male** | **Fw_male** | **CwFw_male** | **Total Males** | ***O. ochracea* collection** | **Category (0 = Fully silent; 0.5 = Low quantity of Singing-capable males; Intermediate/High quantity of Singing-capable males)** |
|---|---|---|---|---|---|---|---|---|
| Hawaii.CL | 20 | 11 | 0 | 4 | 0 | 35 | 6 | 1 |
| Hawaii.UH | 7 | 12 | 2 | 0 | 0 | 21 | 10 | 1 |
| Kauai.CG* | 0 | 9 | 0 | 5 | 9 | 23 | 18 | 0.5 |
| Kauai.WC | 1 | 0 | 0 | 13 | 5 | 19 | 10 | 0.5 |
| Kauai.AS | 0 | 0 | 0 | 21 | 0 | 21 | 18 | 0 |
| Kauai.PK | 14 | 0 | 0 | 5 | 0 | 19 | 8 | 1 |
| Kauai.KCC* | 0 | 0 | 0 | 16 | 1 | 17 | 14 | 0.5 |
| Kauai.PV | 2 | 1 | 0 | 2 | 1 | 6 | 23 | 0.5 |
| Oahu.CC | 2 | 6 | 0 | 15 | 7 | 30 | 9 | 0.5 |
| Oahu.BYU | 2 | 0 | 0 | 8 | 4 | 14 | 11 | 1 |
| Kauai.BI | 1 | 2 | 0 | 2 | 4 | 9 | 12 | 0.5 |
| Kauai.HS | 1 | 0 | 0 | 23 | 0 | 24 | 12 | 0.5 |

**Supplementary Table 2**: Genomic SNPs showing significant association with proportion of singing-capable host males across sites across the Hawaiian archipelago in *LFMM* analysis

| Chromosome | Position | *B* | Adjusted p-value | Distance from nearest gene | Gene class | Genename | Gene ID |
|---|---|---|---|---|---|---|---|
| 1 | 12100553 | 0.466291529 | 0.028413226 | 42761 | Intergenic | Unannotated | ENSEGHG00000003218 |
| 1 | 50888028 | -0.649450864 | 0.038571223 | 0 | Genic | GTF2A1 | ENSEGHG00000001205 |
| 2 | 16669673 | 0.503681033 | 0.017764625 | 27922 | Intergenic | CCDC66 | ENSEGHG00000009961 |
| 2 | 16984343 | 0.41285056 | 0.048324292 | 0 | Genic | PAK5 | ENSEGHG00000006534 |
| 2 | 17774978 | 0.321342801 | 0.046664827 | 46335 | Intergenic | Unannotated | ENSEGHG00000008824 |
| 2 | 25348621 | 0.330628968 | 0.023028903 | 17562 | Intergenic | Unannotated | ENSEGHG00000008088 |
| 2 | 25512221 | 0.330628968 | 0.023028903 | 44406 | Intergenic | Unannotated | ENSEGHG00000004688 |
| 2 | 26989043 | 0.412924959 | 0.017764625 | 7466 | Intergenic | Unannotated | ENSEGHG00000008444 |
| 2 | 27069067 | 0.395463802 | 0.020705672 | 38101 | Intergenic | Unannotated | ENSEGHG00000006501 |
| 2 | 27539143 | 0.40391407 | 0.017764625 | 30684 | Intergenic | NOVA1 | ENSEGHG00000010480 |
| 2 | 27654538 | 0.40391407 | 0.017764625 | 998 | Intergenic | Unannotated | ENSEGHG00000008616 |
| 2 | 27708165 | 0.386452913 | 0.020705672 | 0 | Exonic | Unannotated | ENSEGHG00000008938 |
| 2 | 27728469 | 0.338443746 | 0.035555266 | 0 | Genic | Unannotated | ENSEGHG00000011524 |
| 2 | 27774388 | 0.365377513 | 0.038294072 | 888 | Intergenic | Unannotated | ENSEGHG00000009782 |
| 2 | 27878487 | 0.451485531 | 0.035900519 | 19129 | Intergenic | POU3F2 | ENSEGHG00000011601 |
| 2 | 27967692 | 0.442474134 | 0.020878869 | 0 | Genic | Unannotated | ENSEGHG00000008608 |
| 2 | 29998003 | 0.442515913 | 0.035687273 | 0 | Genic | MSI2 | ENSEGHG00000009793 |
| 2 | 30200480 | 0.338905467 | 0.048324292 | 435 | Intergenic | BLM | ENSEGHG00000006857 |
| 2 | 30796976 | 0.462887141 | 0.023028903 | 1536 | Intergenic | Unannotated | ENSEGHG00000004759 |
| 2 | 30889211 | 0.341060594 | 0.034166994 | 0 | Genic | MITF | ENSEGHG00000008045 |
| 2 | 30924395 | 0.471791047 | 0.029265179 | 0 | Genic | MITF | ENSEGHG00000008045 |
| 2 | 31373007 | 0.410769833 | 0.020705672 | 55387 | Intergenic | Unannotated | ENSEGHG00000010135 |
| 2 | 31378807 | 0.349595786 | 0.035555266 | 61187 | Intergenic | Unannotated | ENSEGHG00000010135 |
| 2 | 31380450 | 0.410769833 | 0.020705672 | 62830 | Intergenic | Unannotated | ENSEGHG00000010135 |
| 2 | 31417220 | 0.40391407 | 0.017764625 | 99600 | Intergenic | Unannotated | ENSEGHG00000010135 |
| 2 | 31494118 | 0.349595786 | 0.035555266 | 73916 | Intergenic | Unannotated | ENSEGHG00000011129 |
| 2 | 31563777 | 0.412924959 | 0.017764625 | 4257 | Intergenic | Unannotated | ENSEGHG00000011129 |
| 2 | 31596070 | 0.341060594 | 0.034166994 | 0 | Genic | Unannotated | ENSEGHG00000011129 |
| 2 | 31786387 | 0.341060594 | 0.034166994 | 0 | Exonic | bco1l | ENSEGHG00000007810 |
| 2 | 31815306 | 0.383836066 | 0.048324292 | 0 | Genic | RRM2B | ENSEGHG00000007972 |
| 2 | 32004163 | 0.412924959 | 0.017764625 | 0 | Genic | Unannotated | ENSEGHG00000010998 |
| 2 | 32044884 | 0.350071483 | 0.035814269 | 0 | Genic | Unannotated | ENSEGHG00000008172 |
| 2 | 32049699 | 0.395463802 | 0.020705672 | 0 | Genic | Unannotated | ENSEGHG00000008172 |
| 2 | 32190767 | 0.44885727 | 0.035687273 | 1560 | Intergenic | Unannotated | ENSEGHG00000007774 |
| 2 | 32668190 | 0.384560834 | 0.036553801 | 9506 | Intergenic | Unannotated | ENSEGHG00000007936 |

| | | | | | | | |
|---|---|---|---|---|---|---|---|
| 2 | 43513377 | 0.666906846 | 0.041128463 | 20649 | Intergenic | Unannotated | ENSEGHG00000009909 |
| 2 | 58284417 | -0.495732583 | 0.038294072 | 1844 | Intergenic | Unannotated | ENSEGHG00000008875 |
| 2 | 58381486 | 0.486943405 | 0.048324292 | 8389 | Intergenic | Unannotated | ENSEGHG00000006706 |
| 2 | 58414464 | -0.586110571 | 0.017764625 | 0 | Genic | Unannotated | ENSEGHG00000005996 |
| 2 | 58497796 | -0.520646499 | 0.017764625 | 0 | Exonic | PRDM5 | ENSEGHG00000008170 |
| 2 | 58510017 | -0.503080151 | 0.042678314 | 5027 | Intergenic | Unannotated | ENSEGHG00000010586 |
| 2 | 58520183 | 0.518975484 | 0.035439356 | 0 | Exonic | Unannotated | ENSEGHG00000011125 |
| 2 | 58561647 | 0.515772201 | 0.017764625 | 0 | Genic | GRIK1 | ENSEGHG00000007865 |
| 2 | 58566560 | -0.512356791 | 0.035687273 | 0 | Genic | Unannotated | ENSEGHG00000009874 |
| 2 | 58574313 | -0.51096372 | 0.035687273 | 6904 | Intergenic | Unannotated | ENSEGHG00000009874 |
| 2 | 58579799 | 0.459426209 | 0.049238174 | 12390 | Intergenic | Unannotated | ENSEGHG00000009874 |
| 2 | 58591705 | -0.504981513 | 0.017764625 | 24296 | Intergenic | Unannotated | ENSEGHG00000009874 |
| 2 | 58681112 | 0.494949064 | 0.020705672 | 53336 | Intergenic | Unannotated | ENSEGHG00000006129 |
| 2 | 58744695 | -0.510741369 | 0.035687273 | 81323 | Intergenic | TLR3 | ENSEGHG00000009430 |
| 2 | 58787587 | -0.520508762 | 0.020705672 | 38431 | Intergenic | TLR3 | ENSEGHG00000009430 |
| 2 | 58804640 | 0.466422547 | 0.020878869 | 21378 | Intergenic | TLR3 | ENSEGHG00000009430 |
| 2 | 58829212 | -0.469689646 | 0.028413226 | 0 | Exonic | TLR3 | ENSEGHG00000009430 |
| 2 | 58835838 | 0.4360392 | 0.049238174 | 4644 | Intergenic | TLR3 | ENSEGHG00000009430 |
| 2 | 58907362 | 0.483334726 | 0.020878869 | 76168 | Intergenic | TLR3 | ENSEGHG00000009430 |
| 2 | 58941143 | -0.49740066 | 0.023028903 | 109949 | Intergenic | TLR3 | ENSEGHG00000009430 |
| 2 | 59029987 | 0.463177361 | 0.035439356 | 126196 | Intergenic | Unannotated | ENSEGHG00000011048 |
| 5 | 12008371 | 0.708916917 | 0.020563143 | 84078 | Intergenic | FKBP8 | ENSEGHG00000001585 |
| 5 | 12099680 | 0.618055782 | 0.017764625 | 0 | Genic | KIF1B | ENSEGHG00000002517 |
| 5 | 19925605 | 0.669778563 | 0.047341268 | 96 | Intergenic | Unannotated | ENSEGHG00000003944 |
| 5 | 20576812 | 0.696493666 | 0.035687273 | 0 | Genic | Unannotated | ENSEGHG00000003363 |
| 5 | 20621916 | 0.715716859 | 0.035814269 | 27075 | Intergenic | Unannotated | ENSEGHG00000003363 |
| 5 | 20698366 | 0.65522943 | 0.035687273 | 29141 | Intergenic | Unannotated | ENSEGHG00000001170 |
| 5 | 20698467 | 0.709677721 | 0.022380845 | 29040 | Intergenic | Unannotated | ENSEGHG00000001170 |
| 5 | 20767146 | 0.746011058 | 0.017764625 | 14597 | Intergenic | SDHB | ENSEGHG00000002681 |
| 5 | 20885498 | 0.759934013 | 0.017764625 | 576 | Intergenic | TEX15 | ENSEGHG00000003964 |
| 5 | 20957234 | 0.895425121 | 0.001361828 | 928 | Intergenic | Unannotated | ENSEGHG00000001543 |
| 5 | 31016229 | -0.776252099 | 0.038294072 | 1028 | Intergenic | SDHB | ENSEGHG00000003685 |

**Supplementary Table 3**: Hardy-Weinberg Equilibrium test on SNPs showing significant association in gene-by-environment scan.

| Chromosome | Position | Observed (HomRef/Het/HomAlt) | Expected (HomRef/Het/HomAlt) | χ2 (HWE test) | P-value |
|---|---|---|---|---|---|
| 1 | 12100553 | 121/30/0 | 122.49/27.02/1.49 | 1.836884 | 0.3631775 |
| 1 | 50888028 | 87/64/0 | 93.78/50.44/6.78 | 10.919 | 0.000347074 |
| 2 | 16669673 | 123/25/3 | 121.59/27.82/1.59 | 1.549438 | 0.1876195 |
| 2 | 16984343 | 127/21/3 | 125.21/24.59/1.21 | 3.212484 | 0.09564116 |
| 2 | 17774978 | 130/18/3 | 127.95/22.09/0.95 | 5.182053 | 0.05056616 |
| 2 | 25348621 | 142/9/0 | 142.13/8.73/0.13 | 0.1424711 | 1 |
| 2 | 25512221 | 142/9/0 | 142.13/8.73/0.13 | 0.1424711 | 1 |
| 2 | 26989043 | 138/13/0 | 138.28/12.44/0.28 | 0.3055399 | 1 |
| 2 | 27069067 | 138/13/0 | 138.28/12.44/0.28 | 0.3055399 | 1 |
| 2 | 27539143 | 139/12/0 | 139.24/11.52/0.24 | 0.2585493 | 1 |
| 2 | 27654538 | 139/12/0 | 139.24/11.52/0.24 | 0.2585493 | 1 |
| 2 | 27708165 | 139/12/0 | 139.24/11.52/0.24 | 0.2585493 | 1 |
| 2 | 27728469 | 141/10/0 | 141.17/9.67/0.17 | 0.177097 | 1 |
| 2 | 27774388 | 139/12/0 | 139.24/11.52/0.24 | 0.2585493 | 1 |
| 2 | 27878487 | 119/32/0 | 120.70/28.61/1.70 | 2.121043 | 0.3754118 |
| 2 | 27967692 | 121/30/0 | 122.49/27.02/1.49 | 1.836884 | 0.3631775 |
| 2 | 29998003 | 123/27/1 | 123.39/26.22/1.39 | 0.135317 | 1 |
| 2 | 30200480 | 140/11/0 | 140.20/10.60/0.20 | 0.2157627 | 1 |
| 2 | 30796976 | 128/23/0 | 128.88/21.25/0.88 | 1.026182 | 1 |
| 2 | 30889211 | 141/10/0 | 141.17/9.67/0.17 | 0.177097 | 1 |
| 2 | 30924395 | 127/24/0 | 127.95/22.09/0.95 | 1.125408 | 0.6002738 |
| 2 | 31373007 | 137/14/0 | 137.32/13.35/0.32 | 0.3568191 | 1 |
| 2 | 31378807 | 140/11/0 | 140.20/10.60/0.20 | 0.2157627 | 1 |
| 2 | 31380450 | 137/14/0 | 137.32/13.35/0.32 | 0.3568191 | 1 |
| 2 | 31417220 | 139/12/0 | 139.24/11.52/0.24 | 0.2585493 | 1 |
| 2 | 31494118 | 140/11/0 | 140.20/10.60/0.20 | 0.2157627 | 1 |
| 2 | 31563777 | 138/13/0 | 138.28/12.44/0.28 | 0.3055399 | 1 |
| 2 | 31596070 | 141/10/0 | 141.17/9.67/0.17 | 0.177097 | 1 |
| 2 | 31786387 | 141/10/0 | 141.17/9.67/0.17 | 0.177097 | 1 |
| 2 | 31815306 | 140/10/1 | 139.24/11.52/0.24 | 2.63836 | 0.2032809 |
| 2 | 32004163 | 138/13/0 | 138.28/12.44/0.28 | 0.3055399 | 1 |
| 2 | 32044884 | 140/11/0 | 140.20/10.60/0.20 | 0.2157627 | 1 |

| | | | | | |
|---|---|---|---|---|---|
| 2 | 32049699 | 138/13/0 | 138.28/12.44/0.28 | 0.3055399 | 1 |
| 2 | 32190767 | 121/30/0 | 122.49/27.02/1.49 | 1.836884 | 0.3631775 |
| 2 | 32668190 | 123/28/0 | 124.30/25.40/1.30 | 1.576855 | 0.6162988 |
| 2 | 43513377 | 1/61/89 | 6.57/49.86/94.57 | 7.541716 | 0.005409223 |
| 2 | 58284417 | 1/27/123 | 1.39/26.22/123.39 | 0.135317 | 1 |
| 2 | 58381486 | 124/26/1 | 124.30/25.40/1.30 | 0.08311985 | 1 |
| 2 | 58414464 | 2/26/123 | 1.49/27.02/122.49 | 0.2151286 | 0.6398659 |
| 2 | 58497796 | 0/29/122 | 1.39/26.22/123.39 | 1.703914 | 0.3630134 |
| 2 | 58510017 | 1/25/125 | 1.21/24.59/125.21 | 0.04279609 | 1 |
| 2 | 58520183 | 124/26/1 | 124.30/25.40/1.30 | 0.08311985 | 1 |
| 2 | 58561647 | 126/25/0 | 127.03/22.93/1.03 | 1.229978 | 0.5998987 |
| 2 | 58566560 | 1/26/124 | 1.30/25.40/124.30 | 0.08311985 | 1 |
| 2 | 58574313 | 1/28/122 | 1.49/27.02/122.49 | 0.1986918 | 1 |
| 2 | 58579799 | 124/27/0 | 125.21/24.59/1.21 | 1.45559 | 0.6080766 |
| 2 | 58591705 | 0/26/125 | 1.12/23.76/126.12 | 1.340002 | 0.6025468 |
| 2 | 58681112 | 127/24/0 | 127.95/22.09/0.95 | 1.125408 | 0.6002738 |
| 2 | 58744695 | 1/27/123 | 1.39/26.22/123.39 | 0.135317 | 1 |
| 2 | 58787587 | 1/25/125 | 1.21/24.59/125.21 | 0.04279609 | 1 |
| 2 | 58804640 | 128/23/0 | 128.88/21.25/0.88 | 1.026182 | 1 |
| 2 | 58829212 | 0/25/126 | 1.03/22.93/127.03 | 1.229978 | 0.5998987 |
| 2 | 58835838 | 126/25/0 | 127.03/22.93/1.03 | 1.229978 | 0.5998987 |
| 2 | 58907362 | 125/26/0 | 126.12/23.76/1.12 | 1.340002 | 0.6025468 |
| 2 | 58941143 | 0/27/124 | 1.21/24.59/125.21 | 1.45559 | 0.6080766 |
| 2 | 59029987 | 127/24/0 | 127.95/22.09/0.95 | 1.125408 | 0.6002738 |
| 5 | 12008371 | 82/67/2 | 88.35/54.31/8.35 | 8.247341 | 0.003093943 |
| 5 | 12099680 | 109/42/0 | 111.92/36.16/2.92 | 3.940296 | 0.07887829 |
| 5 | 19925605 | 55/93/3 | 68.23/66.55/16.23 | 23.86159 | 2.94E-07 |
| 5 | 20576812 | 67/81/3 | 76.53/61.94/12.53 | 14.30388 | 0.000110194 |
| 5 | 20621916 | 77/71/3 | 83.82/57.37/9.82 | 8.52692 | 0.004479958 |
| 5 | 20698366 | 69/81/1 | 79.41/60.19/11.41 | 18.05274 | 3.85E-06 |
| 5 | 20698467 | 64/85/2 | 75.11/62.77/13.11 | 18.93525 | 3.79E-06 |
| 5 | 20767146 | 67/81/3 | 76.53/61.94/12.53 | 14.30388 | 0.000110194 |
| 5 | 20885498 | 62/86/3 | 73.01/63.97/14.01 | 17.90058 | 1.34E-05 |
| 5 | 20957234 | 57/90/4 | 68.90/66.20/15.90 | 19.52004 | 5.18E-06 |
| 5 | 31016229 | 92/54/5 | 93.78/50.44/6.78 | 0.7535076 | 0.4725327 |

**Supplementary Table 4**: Estimation of selection coefficients of two SNPs found within two candidate genes (GRIK1 and TLR3). For each SNP, selection coefficients were inferred for three nested models (1 epoch, 2 epoch and 3 epoch models). Models with the greatest support

| chr | position | epoch | logLR | log10 (p-value) | Epoch1 (start) | Epoch1 (end) | SelectionMLE1 | Epoch2 (start) | Epoch2 (end) | SelectionMLE2 | Epoch3 (start) | Epoch3 (end) | SelectionMLE3 | Gene |
|---|---|---|---|---|---|---|---|---|---|---|---|---|---|---|
| 2 | 58561647 | 1 | 573.0503 | 250.5 | 0 | 10000 | 0.02809 | | | | | | | GRIK1 |
| 2 | 58561647 | 2 | 573.0508 | 248.87 | 0 | 1000 | 0.02804 | 1000 | 10000 | -0.00402 | | | | GRIK1 |
| 2 | 58561647 | 3 | 623.4168 | 269.3 | 0 | 200 | 0.00548 | 200 | 1000 | 0.07567 | 1000 | 10000 | -0.00813 | GRIK1 |
| 2 | 58829212 | 1 | 31.346 | 14.62 | 0 | 10000 | 0.03171 | | | | | | | TLR3 |
| 2 | 58829212 | 2 | 31.346 | 13.61 | 0 | 1000 | 0.03169 | 1000 | 10000 | -0.01584 | | | | TLR3 |
| 2 | 58829212 | 3 | 32.5445 | 13.32 | 0 | 200 | 0.03849 | 200 | 1000 | 0.01267 | 1000 | 10000 | -0.02426 | TLR3 |

**Supplementary Table 6**: Tachinid species used for comparative genomic analyses. Species randomly subsampled from a larger dataset of tachinids with available chromosome-level genomes on NCBI. Number of chromosomes and sex determination taken from genome notes on NCBI from Tree of Life genome curators and is based on synteny to other tachinids. Notes on ecology primarily from Belshaw (1993).

| Species | Genome size ( Mb) | Sex determination system | Size of X chromosome ( Mb) | Number of chromosomes | Host | Notes on ecology. |
|---|---|---|---|---|---|---|
| *Nowickia ferox* | 670.7 | X0 | 15.13 | 5 Autosomes; X chromosome. | Dark Arches moth | Host details from Belshaw (1993). Position in phylogeny is unresolved. One brood (Tschorsnig & Herting, 1994). |
| *Thecocarcelia acutangulata* | 692.9 | X0 | 10.68 | 5 Autosomes; X chromosome. | Lepidoptera | Host details from Dobson (1996) |
| *Tachina grossa* | 936.9 | X0 | 18.03 | 5 Autosomes; X chromosome. | Noctuid moth caterpillars | Bellshaw (1993) |
| *Tachina fera* | 751.7 | X0 | 14.18 | 5 Autosomes; X chromosome. | Noctuid moth caterpillars | Host details from Bellshaw (1993); Mother lays eggs on leaves, larvae then hatch and make their way to host stimulated by vibration. |
| *Epicampocera succincta* | 398.1 | XY | 13.94 | 5 autosomes; X and Y chromosome | *Pieris butterflies (caterpillars)* | Bellshaw (1993) |
| *Gymnosoma rotundatum* | 779.1 | Unknown (X0 or XY) | 17.34 | 5 autosomes X. Unknown whether Y is present. | Shieldbugs | Bellshaw (1993) |
| *Ormia ochracea* | 330.9 | X0 | 9.85 | 5 Autosomes; X chromosomes. | Gryllus and Teleogryllus. | |
| *Cistogaster globosa* | 837.8 | XY | 26.89 | 5 autosomes; X and Y chromosome | Shieldbugs | Raper and Smith (2002) |
| *Tachina lurida* | 899.2 | X0 | 18.71 | 5 Autosomes; X chromosome. | Noctuid moth caterpillars | Bellshaw (1993) |
| *Phasia obesa* | 876.8 | X0 | 24.2 | 5 Autosomes; X chromosome. | *Lygus* spp. (tarnished plant bugs) | Rämert, B., Hellqvist, S., & Petersen, M. K. (2005). |
